# Assessing the Molecular Validity of Spontaneous Lupus Mouse Models and Its Implication for Human Studies

**DOI:** 10.1101/2025.05.15.654304

**Authors:** María Rivas-Torrubia, María Morell, Zuzanna Makowska, Jorge Kageyama, Anne Bosshard, Julius Lindblom, Maria Orietta Borghi, Eleonore Bettacchioli, PRECISESADS flow cytometry consortium, PRECISESADS clinical consortium, Ioannis Parodis, Lorenzo Beretta, Concepción Marañón, Jacques-Olivier Pers, Ralf Lesche, Fiona McDonald, Marta E. Alarcón-Riquelme, Guillermo Barturen

## Abstract

**Background:** Systemic lupus erythematosus (SLE) is a complex autoimmune disease characterized by a loss of self-tolerance, causing inflammation and tissue damage in multiple organs. Animal models have advanced our understanding of SLE’s molecular basis, but the FDA’s recent elimination of animal testing requirements for drug approval has raised concerns about their validity, prompting a reevaluation of their role in basic research, especially for heterogeneous diseases like SLE.

**Methods:** Four different spontaneous SLE mouse models were studied: MRL*^lpr/lpr^*, NZB/W, BXSB.*Yaa*, and Tlr7.Tg6. Transcriptome sequencing from blood, spleen, and kidney, flow cytometry from the spleen, and cytokines and autoantibody measurement in plasma were performed at four time points. Similar molecular data from human SLE patients was used for the integration.

**Results:** The study identified specific molecular pathways driving the phenotype in each mouse model and established optimal time points for future experimental designs. By comparing these pathways to human SLE, the most similar ones and their relationship with disease activity were identified, providing crucial insight into translational relevance.

Importantly, disease severity across models was linked to the extent and timing of molecular dysregulations. As expected, MRL*^lpr/lpr^* showed the most aggressive phenotype with early immune activation and apoptosis dysregulation, while Tlr7.Tg6 presented late-onset signatures associated with interferon and inflammation. Shared molecular features with human SLE included interferon responses, T and B cell depletion, and neutrophil activation. Integration analysis revealed distinct yet overlapping immune pathways between models and species, with some signatures such as age-associated B cells and double-negative memory T cells being model-specific but potentially relevant to early disease processes.

**Conclusions:** These findings build a valuable framework for future SLE research, reinforcing the utility of mouse models in studying specific molecular pathways related to human SLE pathogenesis and heterogeneity. The integration of longitudinal mouse data with human transcriptomes highlights the models that best recapitulate key aspects of human disease, offering guidance for the study of specific immunopathological mechanisms or therapeutic targets.

## BACKGROUND

The mouse has been a key model organism for studying human diseases, aiding in the understanding of molecular dysregulation in human phenotypes. However, concerns about the reproducibility and translatability of mouse model results to humans, particularly in preclinical trials, have arisen. Recently, the Food and Drug Administration (FDA) was authorized to proceed with human trials without requiring drug safety and efficacy tests in animals (1), questioning the role of mouse models in basic research and prompting a review of their utility in mimicking human diseases.

The animal modeling of complex diseases, such as systemic lupus erythematosus (SLE), is particularly challenging due to diseases’ clinical and molecular heterogeneity (2,3). Recent publications have shown that different subgroups of clinically diagnosed SLE patients show different molecular pathways dysregulations during the course of the disease (4,5), some of which are shared with other systemic autoimmune diseases (6). This molecular heterogeneity might partially explain the failure of some randomized clinical trials of treatments commonly used for other diseases, or even for SLE in the clinical practice (7).

Numerous mouse models exist for studying SLE, though none fully replicate the entire disease’s clinical spectrum. Each model exhibits human-like phenotypes and offers specific characteristics of preclinical interest (8), mimicking the molecular heterogeneity found in SLE patients. Two types of models can be found: induced and spontaneous. Induced models are useful for the study of environmental factors contributing to the pathogenesis of the disease, while spontaneous models result from genetic mutations or duplications making them adequate for exploring the genetic contributions to SLE pathogenesis. It is crucial to consider molecular heterogeneity during animal model selection for the success of treatment trials or the study of the implication of specific SLE-related molecular pathways in the course and pathophenotype of the disease (9).

Four spontaneous mouse models were studied: BXSB.*Yaa*, Tlr7.Tg6, MRL*^lpr/lpr^* and NZB/W. BXSB.*Yaa* and Tlr7.Tg6 involve *Tlr7* gene duplication, in the case of BXSB.*Yaa* the entire Y-autoimmune accelerator locus was translocated in males from the X to the Y chromosome. The *Tlr7* gene and the *Yaa* locus regulate the activation of the type I interferon pathway by ribonucleic acid complexes, a critical pathway in the pathogenesis of SLE. Despite sharing *Tlr7* gene duplication, they differ in cell population expansion, inflammatory pathways, autoantibody positivity, and disease progression (10,11). The MRL*^lpr/lpr^* model, driven by the *lpr* mutation in the *Fas* gene, leads to malfunctioning of immune cell apoptosis resulting in the accumulation of CD4-CD8 double-negative T cells and the induction of the disease due to the amplification of susceptibility genes. It is a model with rapid and severe development compared to other spontaneous models (11). Lastly, the NZB/W model is a F1 hybrid between the New Zealand Black (NZB) and the New Zealand White (NZW) strains. It is characterized by early elevated antinuclear antibodies (ANAs) levels and other diverse autoantibodies which lead to immune complex-mediated nephritis and a fatal outcome (11).

This study profiled similar molecular data from different omic layers in mouse and human patients from the PRECISESADS SLE cohort (6). Considering the heterogeneity of SLE patients and the diversity of the available SLE mouse models, we reasoned that studying SLE patients and mouse models together at various molecular layers and across different tissues was crucial. This approach not only characterized the mouse models in-depth but also defined their molecular similarities and differences with the human disease. Using novel molecular data integration factorization methodology, we aimed to establish a framework for the future design of preclinical studies in SLE, highlighting the models exhibiting molecular signatures similar to those associated with human disease activity and the time required for the models to develop human-like molecular signatures.

## METHODS

### Cohorts and Study Design

#### Human cohort

Whole blood samples from 342 SLE patients and 497 healthy controls were obtained from the PRECISESADS cohort (Table S1). Recruitment criteria and detailed information regarding the sampling and the molecular profiling can be found in Barturen et al. (6).

#### Mouse models

Whole blood, spleen, and kidney were sampled at four time points for four SLE spontaneous mouse models (MRL*^lpr/lpr^*, NZB/W F1, BXSB.*Yaa*, and Tlr7.Tg6), as well as for their respective genetic controls (MRL/J, NZW, BXSB, C57BL/6). Since the severity and the kinetics of the development of the SLE phenotype in each model are different, different time points were used to try to synchronize the development of the phenotypes between models. In MRL*^lpr/lpr^*, samples were taken at weeks 6, 12, 14 and 16; in NZB/W, weeks 6, 12, 18 and 28; in BXSB.*Yaa*, weeks 6, 12, 16 and 20; and in Tlr7.Tg6, weeks 6, 12, 16 and 28. The different weeks will be referred to as pseudotimes 1,2, 3 and 4.

#### Sex as a biological variable

The human dataset includes both males and females. The MRL^lpr/lpr^ and NZB/W F1 mice and their respective controls were females, while the BXSB.*Yaa* and Tlr7.Tg6 mice were males. The development of the disease in these spontaneous models is driven by genetic modifications that lead to a SLE-like phenotype. Therefore, the molecular features associated with disease development should be studied in comparison to their respective genetic background, including sex, to ensure that observed changes are not influenced by sex differences. The male models used in this study involve either the translocation of Y-autoimmune accelerator locus from X chromosome to Y, or the duplication of *Tlr7*, which mimics Tlr7 overexpression as may occur in females due to X chromosome inactivation escape. As a result, these models are not directly comparable to human sex-related differences in SLE. Consequently, the disease phenotypes are driven by genetic modifications rather than biological sex.

#### Mouse sampling of blood, kidney and spleen

At each time point, 10 animals from each SLE model and 5 controls were sacrificed (Table S2). Mice were anesthetized by an intraperitoneal injection of 100 ug/g body weight ketamine and 16ug/g body weight xylazine. After the anesthesia, blood was collected through cardiac puncture and transferred into ethylenediaminetetraacetic acid (EDTA)-coated tubes (Vacutainer, Beckman Dickinson). The animals were perfused with cold phosphate-buffered saline (PBS), and the spleen and both kidneys were collected into cold PBS. Whole anticoagulated blood was centrifuged for 5min at 500g and 4□C to separate the plasma and stored at -80□C. The cell pellets were resuspended in 10 ml red blood cell lysis buffer (RBCL; 155 mM NH4Cl, 12 mM NaHCO3, 0.1 mM EDTA) and incubated for 5 minutes at room temperature. Leukocyte pellets were washed with cold MACS buffer (Miltenyi) and resuspended in the same volume of MACS buffer as the initial blood volume. 200 ul of the cell suspension was centrifuged for 1 minute at 10000 g, the cell pellet was immediately frozen on dry ice and stored at -80□C until RNA extraction. A fragment of the spleen was homogenized. Red cells were lysed as before. Splenocytes were resuspended in MACS buffer, counted and analyzed by flow cytometry.

#### Transcriptome profiling

RNA was extracted from snap-frozen white blood cell pellets, and homogenized spleen and kidney pieces using the RNeasy96 kit (Qiagen). For library synthesis, 400 ng of total RNA from the solid tissues and 100 ng of total RNA from white blood cells were used with the TruSeq Stranded mRNA HT kit (Illumina). The libraries were quantified using qPCR with the PerfeCTa NGS kit (Quantart Biosciences), and equimolar amounts of samples were pooled. Pooled samples were clustered on a high-output flow cell using the HiSeq SR Cluster kit v4 and the cBot instrument (Illumina). Subsequently, 50 cycles of single-read sequencing were performed on a HiSeq2500 instrument using a HiSeq SBS kit v4 (Illumina). On average, libraries were sequenced at 11.3 ± 2.95 million reads.

Reads were quality (Q<20) and adapter based trimmed at the 3’ end using the cutadapt software (12). Reads below 25 nucleotides were discarded from the analysis. The resulting reads were aligned against the GRCm38 mouse assembly using the STAR program (13) with default parameters. Gene expression quantification was performed by the RSEM program (14) using the GENCODE annotation (15). Genes with at least 10 reads in 5 samples were selected for further analysis: 18936 in spleen, 16225 in blood and 18491 in kidney.

#### Flow cytometry profiling

Spleen samples were analyzed by flow cytometry using 3 multi-color panels. Panel 1 included B cells (B220+), and other subpopulations with specific markers CD93, B220, IgD, IgM CD21, CD23, CD5. Panel2 included T cells (CD5) and other populations defined by CD4, CD8, CD25, CD49b, CD44, CD62L. Panel 3 included plasma cells (PCs) (CD138), granulocytes (CD11b), neutrophils/eosinophils (CD11b, Gr-1), monocytes (CD11b, CD115), pDCs (CD11c, SiglecH) and cDCs (CD11c, SiglecH, CD8a, CD11b). The gating strategy and definition of all populations can be consulted in Fig. S1 and antibody commercial references are included in Table S3. Samples were stained with the antibody cocktails in a FACSVerse (BXSB.*Yaa* and Tlr7.Tg6) and a FACSCanto II (NZB/W and MRL^lpr/lpr^). The two cytometers were aligned using compensation beads at the beginning of the study, and the fluorescence signals were set up by using 8-peak beads, as described before (16). The compensation matrices and cell proportion quantification were calculated using the FlowJo.v10.7 program (17).

#### Serology profiling

Serum cytokines and autoantibodies were quantified using the Luminex technology. The panels were defined based on the cytokines and autoantibodies associated with different systemic autoimmune diseases in humans. Cytokines were analyzed using the Cytokine & Chemokine 36-plex Mouse ProcartaPlex Panel 1 Kit (Thermo Fisher). It included several interleukines (IL-10, IL-3, IL-1ß, IL-2, IL-4, IL-5, IL-6, IL-22, IL-9, IL12p70, IL-13, IL-27, IL-23, IL-17A, IL-15/IL-15R, 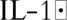, IL-28, IL-18, IL-31), interferons 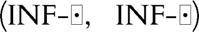, monocyte-colony stimulating factor (M-CSF) granulocyte and monocyte colony stimulating factor (GM-CSF), granulocyte colony stimulating factor (G-CSF), and other cytokines such as leukemia inhibitory factor (LIF), interferon gamma-induced protein 10 (IP-10), growth-regulated oncogene-alpha (GRO-α, also known as CXCL-1), RANTES (regulated on activation, normal T cell expressed and secreted), tumor necrosis factor-alpha (TNF-α), macrophage inflammatory proteins (MIP-1α, MIP-1β, MIP-2), monocyte chemoattractant proteins (MCP-1, MCP-3), epithelial-derived neutrophil-activating peptide 78 (ENA-78), and eotaxin. The auto-antibodies quantified were: anti-histone, anti-Jo1, anti-centromere (anti-CENP-B), anti-double-stranded DNA (anti-dsDNA), anti-Sjögren’s-syndrome-related antigen A (anti-SSA/La), anti-Sjögren’s-syndrome-related antigen B (anti-SSB/Ro), anti-ribonucleoprotein (anti-RNP), and anti-Smith (anti-Sm) and were quantified by adapting the Athena ANA-II Plus Test System kit (Zeus Scientific), using a biotinylated anti-mouse-immunoglobulin G (IgG) antibody as a secondary detection antibody.

### Statistical analysis

#### Data values normalization

All molecular layers were transformed to normalize their distributions. Gene expression values were normalized using the varianceStabilizingTransformation function from the DESeq2 R package (18). Cell proportions, cytokines, and autoantibody values were log10(x+1) transformed.

#### Transcriptional module annotation

Mouse gene expression values were summarized using the GSVA R package (19) into a human immunological module set (20). The homologene database from NCBI (21) was utilized to find homologous genes between human and mouse. In total, 7,892 (69%) homologs were found between mouse and human out of the 11,465 genes annotated in the modules. Modules with less than 10 homologs were discarded.

#### Longitudinal differential expression analysis

Linear regression analyses with an interaction term between group and pseudotime were conducted to identify differential longitudinal molecular entities. The model is represented as follows: Y = β_0_ + β_1_X_1_ + β_2_X_2_ + β_3_X_1_X_2_ + ε, where Y represents the molecular feature quantification, X_1_ represents the mouse groups and X_2_ represents the pseudotime values (1–4). Interaction term analysis identifies the association between molecular entities, mouse model changes and pseudotime. In order to discard significant values due to single point changes, the linear relationship between molecular quantification and pseudotime, solely in the SLE model was conducted as: Y = β_0_ + β_1_X_2_ + ε. Overall, most of the significant molecular entities for the interaction term were also significantly associated with the pseudotime (FDR < 0.05): 294 out of 386 transcriptome modules, 7 out of 7 autoantibodies, 30 out of 31 cytokines, and 20 out of 21 cell type proportions. Significant entities for both linear models were selected for further analyses, indicating molecular differences that either increase or decrease over time.

#### Time point cross-sectional analysis

Individual time point analyses were conducted by means of linear regression models (FDR < 0.05), change direction was confirmed with the longitudinal analysis. Longitudinal molecular differential features were classified as early or late based on the time point analyses. Molecular features with significant differences in 3 or 4 time points were classified as early dysregulations, while late dysregulations were defined as molecular features with significant differences only at pseudo-times 3 and/or 4. Gene functionalities at each pseudo-time were assessed by Gene Set Enrichment Analysis (GSEA) implemented in clusterProfiler R package (22) using MgSigDB hallmark gene set (23).

#### Molecular layer integration

Molecular integration was performed with MEFISTO (24), which based on factor decomposition of the multiple views, summarizes relationships that change gradually over time. SLE patients were classified into five categories based on their SLEDAI-2K: no activity (SLEDAI-2K=0), mild (1 to 5), moderate (6 to 10), high (11–19) and very high (>=20). These categories were based on a previous publication (25,26) and they were assumed to be comparable in terms of severity of the disease with the pseudo-times selected for each mouse model (1 low, 2 moderate, 3 high and 4 very high). All autoantibodies, cytokines and cell populations were included in the analysis for both mouse and human. Transcriptional modules were preselected based on differential analysis between SLE patients and healthy controls (FDR < 0.05) including sex, age, sequencing batch and RNA integrity number as covariates (177 modules were selected). MEFISTO was run using default parameters.

## RESULTS

### Mouse model sampling alignment with disease severity

To investigate the molecular relationships between mouse models and humans, multiple layers of molecular data were profiled from different biological sources and aligned as much as possible between species. Four time points were selected prior to reaching the 50% mortality rate of each model (Fig. 1A), with the aim of capturing the molecular initiation of disease phenotypes. The early time points were chosen to align with the mild-to-moderate disease activity observed in the human SLE PRECISESADS cohort (average SLEDAI-2K: 5.94 ± 5.27).

**Fig. 1:**
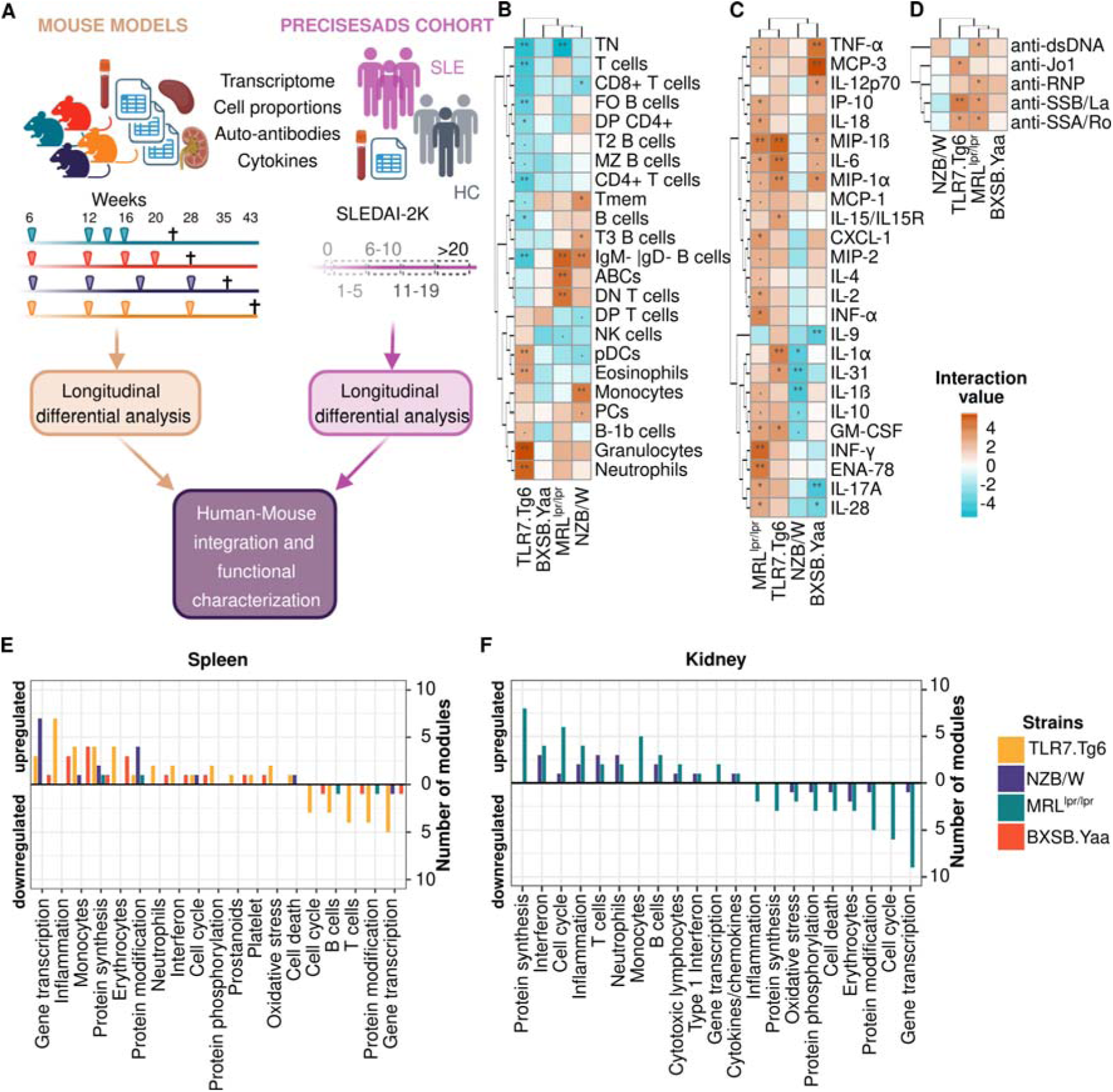
Molecular information shows differences between mouse models. **(A)** Analysis workflow: Different analyses were performed to investigate molecular differences between mice and humans. 50% mortality rates are represented as ✝. Longitudinal significant cell proportions **(B)**, cytokine levels **(C)**, and autoantibody levels **(D)** across mouse models are represented. Interaction term values with at least one comparison below 0.1 FDR are shown. Mouse models and features were grouped by hierarchical clustering. Interaction term significant values were indicated as follows: . FDR < 0.1, * FDR < 0.05, ** FDR < 0.01. The number of longitudinal differential transcriptional immune modules is shown (interaction term FDR < 0.05) for the spleen **(E)** and the kidney **(F)** across the different mouse models. Significant modules were grouped by related molecular functions and sorted by the number and direction of change (from upregulation to downregulation in mouse models compared to their genetic controls). Mouse models are color-coded as follows: yellow (Tlr7.Tg6), purple (NZB/W), green (MRL*^lpr/lpr^*), and red (BXSB.*Yaa*). Modules without defined molecular functions (TBD) were excluded from the plot.

Additionally, time points were selected to align mouse model disease severity, particularly kidney involvement. In the selected time points, the MRL*^lpr/lpr^*, NZB/W and BXSB.*Yaa* models exhibited elevated urine albumin-to-creatinine ratios (ACR) at later time points, indicating kidney dysfunction (Fig. S2A). This increasing trend was consistent with the rising of serum creatinine levels and the presence of abnormal urine creatinine ratios (>0.2) observed in human patients grouped by SLEDAI (Fig. S2B-C). The Tlr7.Tg6 model, known to exhibit the slowest disease progression among the four selected models, did not show abnormal ACR levels even at later time points. Nevertheless, histological analysis of kidney tissue at week 28 revealed evidence of tissue damage, despite the absence of noticeable changes in ACR (Data not shown).

### Dysregulation of specific spleen cell populations characterizes each of the different mouse models

Linear regression models with interaction terms were used to assess the association between cell-type expansions and time-based phenotype development (see Methods). Significant depletion of most lymphoid cell proportions was observed in the Tlr7.Tg6 model, including B and T cell subsets (Fig. 1B). The most notable depletion was that of naïve T cells (TN), a feature common to all models (Fig. 1B) but at different times (Fig. S3A). Specific expansions in T cells, such as memory (Tmem) and double-negative T cells (DN T cells), were seen in the MRL*^lpr/lpr^* and NZB/W models (Fig. 1B and Fig. S3A). In B cells, the MRL*^lpr/lpr^* and NZB/W models showed significant overrepresentation of class-switched B cells (IgM-IgD-B cells in Fig. 1B and Fig. S3A), while age-associated B cells (ABCs) were observed specifically in the MRL*^lpr/lpr^* model (Fig. 1B and Fig. S3A). Time point analyses revealed additional expansions: early expansion of plasma cells (PCs) and marginal zone B cells (MZ B cells) in the MRL*^lpr/lpr^* and NZB/W models, respectively, and late expansion of transitional T3 B cells (T3 B cells) in the NZB/W model (Fig. 1B and Fig. S3A).

The Tlr7.Tg6 and NZB/W models showed the largest myeloid compartment expansion, but through different lineages (Fig. 1B). Eosinophils and plasmacytoid dendritic cells (pDCs) expanded exclusively in the Tlr7.Tg6 model. Notably, pDC expansion was not found in the closely akin BXSB.*Yaa* model (Fig. 1B). Monocytes expanded in the NZB/W model, while neutrophils were expanded in the MRL*^lpr/lpr^* and Tlr7.Tg6 models (Fig. 1B), indicating active inflammation via different pathways. Neutrophils and monocytes changes were not significant in Tlr7.Tg6 and NZB/W time point analyses, respectively, suggesting latter expansions compared to the MRL*^lpr/lpr^* model (Fig. S3A).

### Serum cytokine levels related mostly with cell type expansion in the spleen and revealed an important contribution of macrophages in the BXSB.*Yaa* model

Serum cytokine levels were measured and the panel was selected based on human autoimmunity association. In order to identify cytokines changing over time, interaction terms within linear regression models were used as in the previous analysis.

Overall, the MRL*^lpr/lpr^* model exhibited the highest cytokine production, with significant upregulation of most analyzed cytokines. Particularly, the overexpression of TNF-α, IL-4, IL-2, and IL-17A (Fig. 1C) which may be related with the expansion observed of DN T cells (27,28). This model also showed the highest increase in type I, II (IFN-α and IFN-γ) and III (IFN-λ, IL-28) interferons (Fig. 1C). The timing of cytokine overproduction varied, with early IFN-γ and IP-10, followed by IFN-α, and late IFN-λ (Fig. S3B). Another prominent cytokine and chemokine group may involve neutrophil production and activation, marked by GM-CSF, ENA78, MIP-1α and MIP-1β overproduction. These last two cytokines are also known to induce the release of IL-1, IL-6, and TNF-α (29), all of them overproduced in the model (Fig. 1C).

The Tlr7.Tg6 model also showed significant cytokine overproduction, including GM-CSF, MIP-1α, MIP-1β, and IL-6 (Fig. 1C), but at later stages than the MRL*^lpr/lpr^*model (Fig. S3B). The higher cytokine levels in the Tlr7.Tg6 model correlated with extensive neutrophil expansion as compared to MRL*^lpr/lpr^* (Fig. 1B). The model also showed an eosinophil-specific expansion, potentially related with MIP-1α and MIP-1β overproduction, which are known to activate eosinophil and other myeloid populations (30). Additionally, IL-31 was also overproduction and potentially associated with eosinophil and dendritic cell activity (31,32) (Fig. 1C).

MCP-3 is a cytokine involved in the regulation of macrophage function and monocyte recruitment (33), while IL-12 (IL-12p70) and TNF-α are important pro-inflammatory cytokines produced by macrophages in the targeted tissues (34). All these were highly upregulated in the BXSB.*Yaa* serum, while IL-9, a known anti-inflammatory cytokine for macrophages (35), was downregulated (Fig. 1C). These results suggest a central role of macrophages in the BXSB.*Yaa* model phenotype. A shared neutrophil/macrophage-related cytokine core (MCP-3, MIP-1α, MIP-1β, IL-6, and TNF-α) was identified between MRL*^lpr/lpr^* and BXSB.*Yaa* models (Fig. 1C). Interestingly, the BXSB.*Yaa* model showed earlier overproduction of MIP-1α, MIP-1β, and TNF-α, while MCP-3 and IL-6 appeared later, suggesting different inflammatory initiation pathways and should be further studied in relation to human SLE pathogenesis.

The NZB/W model displayed a different cytokine pattern compared with the other models. Key proinflammatory cytokines, such as IL-1α and IL-1β, are downregulated overtime in the model (Fig. 1C). This distinct cytokine profile indicates a different autoimmune process in the NZB/W model, impacting its phenotype development and progression.

### Autoantibody specificities and timings differentiate SLE mouse models

Autoantibodies associated with human autoimmunity were selected and measured in the serum. The MRL*^lpr/lpr^* and NZB/W models exhibited earlier overproduction of autoantibodies compared to the BXSB.*Yaa* and Tlr7.Tg6 models. The BXSB.*Yaa* model had the fewest autoantibody specificities (Fig. S3C), while the NZB/W model had the most, excluding anti-RNP antibodies (Fig. 1D and Fig. S3C). However, longitudinal analysis did not detect changes in the NZB/W model, likely due to slower autoantibody production from its NZW genetic background, leading to smaller differences over time.

Classic SLE-associated anti-dsDNA autoantibodies were overproduced in all models except Tlr7.Tg6 model (Fig. 1D), occurring earlier in the MRL*^lpr/lpr^* and NZB/W models than in the BXSB.*Yaa* model. Anti-SSA/Ro and anti-SSB/La autoantibodies were highly upregulated in the Tlr7.Tg6 and MRL*^lpr/lpr^* models (Fig. 1D), consistent with high interferon production (Fig. 1C). However, these differences were not apparent in the time point analysis for the MRL*^lpr/lpr^*model, possibly due to late overproduction (Fig. S3C). The NZB/W model also showed overproduction of anti-SSA/Ro and anti-SSB/La antibodies, but was not associated with the interferon production (Fig. 1C).

Anti-Smith (anti-Sm), anti-centromere B, and anti-histone autoantibodies were significantly upregulated early in the MRL*^lpr/lpr^* model (Fig. S3C), but were not significant in the longitudinal analysis (Fig. 1D). Anti-Jo1 autoantibodies were overproduced in both the NZB/W and Tlr7.Tg6 models at different time points (Fig. S3C).

### *Tlr7*-related models showed greater dysregulation of the spleen transcriptome dysregulation

The transcriptome analysis revealed molecular dysregulation in the spleen and kidney across different mouse models. Specifically, the spleen showed significant dysregulation in the Tlr7.Tg6 and the BXSB.*Yaa* models (Fig. 1E and Fig. S4A). In contrast, few or no differentially expressed modules were observed in blood (Fig. S4B), supporting the use of the spleen to study immune system dysregulation in mice models (similar findings were observed at the individual gene level, Additional Files 1-4).

Transcriptional changes in the spleen aligned with the molecular mechanisms from other data layers. In the Tlr7.Tg6 model, interferon, inflammatory, and neutrophil-related modules were upregulated, while lymphoid modules were downregulated (Fig. 1E and Fig. S4A), consistent with spleen cell proportions and cytokine profiles (Fig. 1B-C). The BXSB.*Yaa* model showed upregulation of monocyte and inflammatory modules in the spleen (Fig. 1E and Fig. S4A). Although monocyte module dysregulation was not reflected as cell proportion expansion, cytokine profiles from this model indicated a major role for macrophages, potentially differentiated from mobilized and activated monocytes (Fig. 1C).

### Interferon-related transcriptome dysregulation was primarily observed in the kidneys of MRL*^lpr/lpr^* and NZB/W mice

The kidney transcriptome of both MRL*^lpr/lpr^* and NZB/W models displayed a significant upregulation of interferon-related modules (Fig. 1F and S4C). This upregulation was evident in the MRL*^lpr/lpr^* serum cytokines but not in the NZB/W model (Fig. 1C), making this interferon signature specific to the kidney. Additionally, both models showed an upregulation of neutrophil and lymphoid cell compartments, indicating increased immune cell infiltration in the kidney (Fig. 1F).

### Correlation of blood and spleen transcriptomes validates blood as surrogate for spleen immune monitoring

To investigate the potential sharing of immunological signatures across different tissues, Pearson correlation coefficients between tissues were calculated for all the immunological transcriptional modules for each mouse model (Fig. 2A-C). The analysis revealed a noteworthy correlation between spleen and blood immunological modules across all models, with predominantly positive correlation values for the differentially expressed modules (Fig. 2A and Fig. S5A). However, comparisons between spleen/kidney or blood/kidney modules showed much weaker correlations (Fig. 2B-C). Despite this overall lack of correlation, specific molecular signatures were highly correlated between blood and tissues in the different mouse models. For example, a myeloid inflammatory signature, characterized by monocytes and neutrophils showed significant correlations between spleen and kidney in the BXSB.*Yaa* and MRL*^lpr/lpr^* models (Fig. S5B). Similarly, an interferon signature was observed between blood and kidney in the Tlr7.Tg6 and MRL*^lpr/lpr^* models (Fig. S5C).

**Fig. 2:**
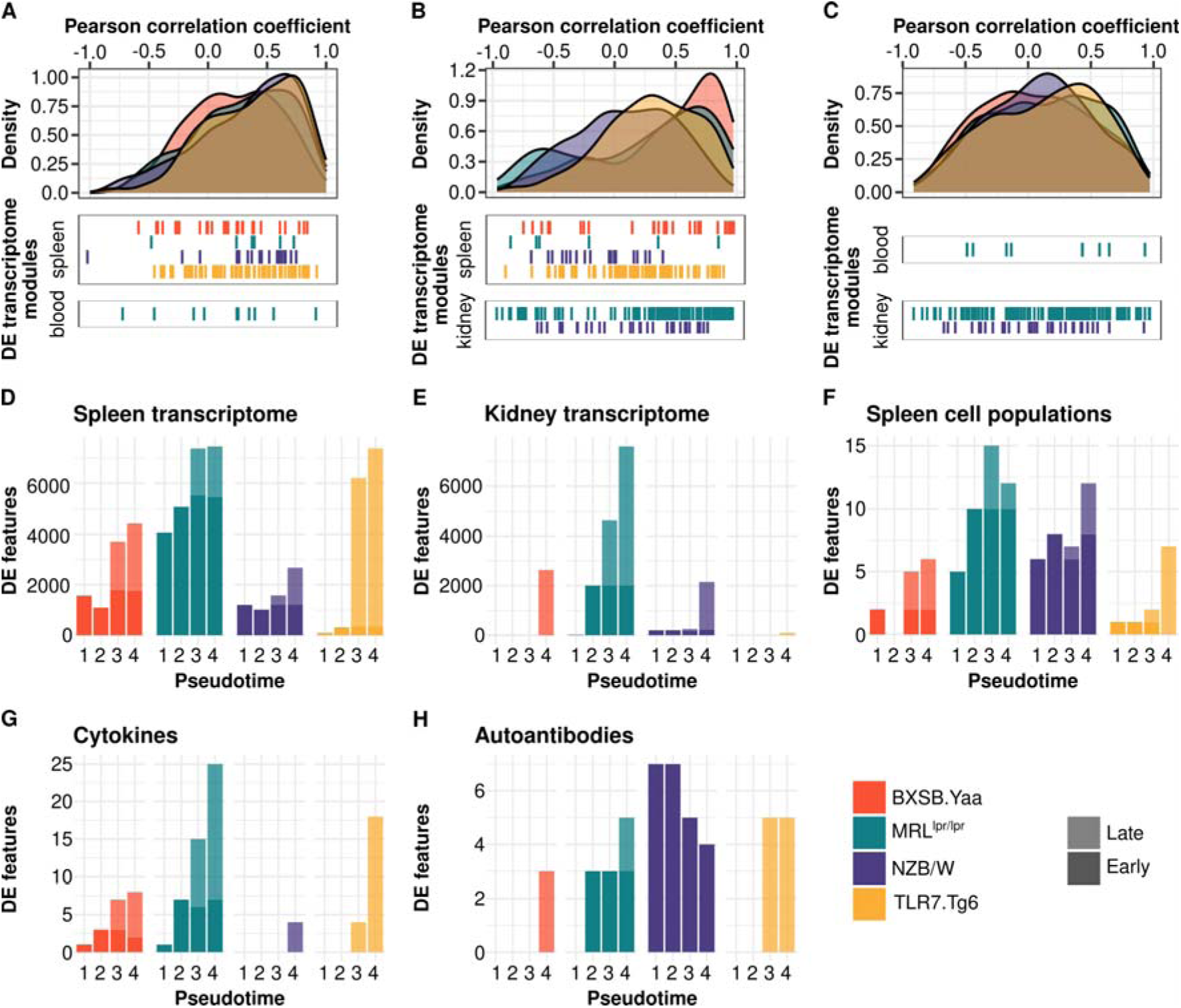
Transcriptome correlations and time point analyses reveal different dysregulation timings across molecular layers and tissues. Density plots for Pearson correlation coefficients are presented for all transcriptional modules by biological sample pairs: spleen-blood **(A)**, spleen-kidney **(B)**, and blood-kidney **(C)**. Significant differentially expressed modules (interaction term FDR < 0.05) are depicted per biological sample and mouse model below the density plots. Bar plots display the cross-sectional analysis at different time points for the spleen transcriptome **(D)**, kidney transcriptome **(E)**, spleen cell proportions **(F)**, cytokines **(G)**, and autoantibodies **(H)**. The differential features per each time point (FDR < 0.05) were classified as early or late. Mouse models are color-coded as follows: yellow (Tlr7.Tg6), purple (NZB/W), green (MRL*^lpr/lpr^*), and red (BXSB.*Yaa*).

### Timing of transcriptome and cytokine dysregulation associated with severity and disease progression

Next, we investigated the relationship between molecular dysregulation and disease severity and/or development of disease in the mouse models. Molecular entities were categorized as early or late based on initial time point when dysregulation was observed and its continuity over time (see Methods).

Gene expression analysis across all mouse models showed that molecular dysregulations in the spleen preceded changes in the kidney (Fig. 2D-E). These changes were earlier and stronger in the MRL*^lpr/lpr^* and BXSB.*Yaa* models compared to NZB/W and Tlr7.Tg6 models. Splenic dysregulation was associated with changes in cell proportions (Fig. 2F) and synchronized with cytokine dysregulation (Fig. 2G). This suggests that immune changes in the spleen may drive subsequent effects on target organs, potentially contributing to the cell infiltration of the tissues. Furthermore, the early molecular changes in the spleen compared to the kidney may in part explain the lower correlations of immune signatures between these organs (Fig. 2B).

Functional pathway enrichment analysis from the first time point revealed dysregulation of immune gene sets in the spleen of all models (Fig. S6A). However, kidney dysregulation in less aggressive NZB/W and Tlr7.Tg6 models, was less significant and/or appeared only at the latest time points (Fig. S6B). Similar patterns were observed for other immune-related gene sets, such as IL2-STAT5 signaling, TNF-α signaling via NFKB, and KRAS signaling (Fig. S6A-B). In the same line, the apoptosis gene set, indicating initiation of tissue damage, was dysregulated earlier in the most aggressive MRL*^lpr/lpr^* and BXSB.*Yaa* models (Fig. S6B). This highlights the association between phenotype timing and the transcriptome dysregulation in the different tissues.

Autoantibody overproduction preceded or became significant at the time points when kidney transcriptome dysregulations became evident (Fig. 2E and Fig. 2H), suggesting a temporal relationship between autoantibody development and molecular changes in the kidney.

### Human SLE molecular signatures were shared between patients and specific mouse models

Mouse-centered analyses revealed molecular signatures potentially driving and/or triggering the phenotype. To investigate whether similar molecular dysregulations underlie the phenotypes in mouse models and in humans, we conducted a disease severity-oriented integration using factor analysis (24).

Severity, or time in mouse models, has been our leading factor for identifying molecular dysregulation in previous analyses. However, SLE in humans does not progress linearly but in waves of flares and remissions, making time since diagnosis as a severity measure useless. In fact, no significant associations were observed between disease duration and molecular features (Additional file 5). The SLEDAI is the most used disease activity index in clinical practice, and it has been previously used to correlate molecular information with successful results (4,5). Indeed, significant associations were found between the SLEDAI-2K and well-known molecular features in our SLE cohort (Fig. S7). Therefore, for integration we defined disease severity in the mouse models as pseudotimes and categorized SLEDAI groups in humans (see Methods).

The integration model resulted in fifteen factors from which four (1, 2, 3, and 5) were selected based on their smoothness (dependency on severity) and a minimum explained variance (Fig. 3A). These four factors were independent (Fig. S8A) and accounted for 68.07±9.85% of the overall observed variance. All molecular layers were represented in the factors, but humans had a diminished cytokine-related signal due to the limited number of cytokines profiled (Fig. 3A). Factors 2 and 3 showed significant trends in humans (Fig. 3B, linear regression p-value < 0.05), while factors 1 and 5 were mouse-specific (Fig. S8B).

**Fig. 3:**
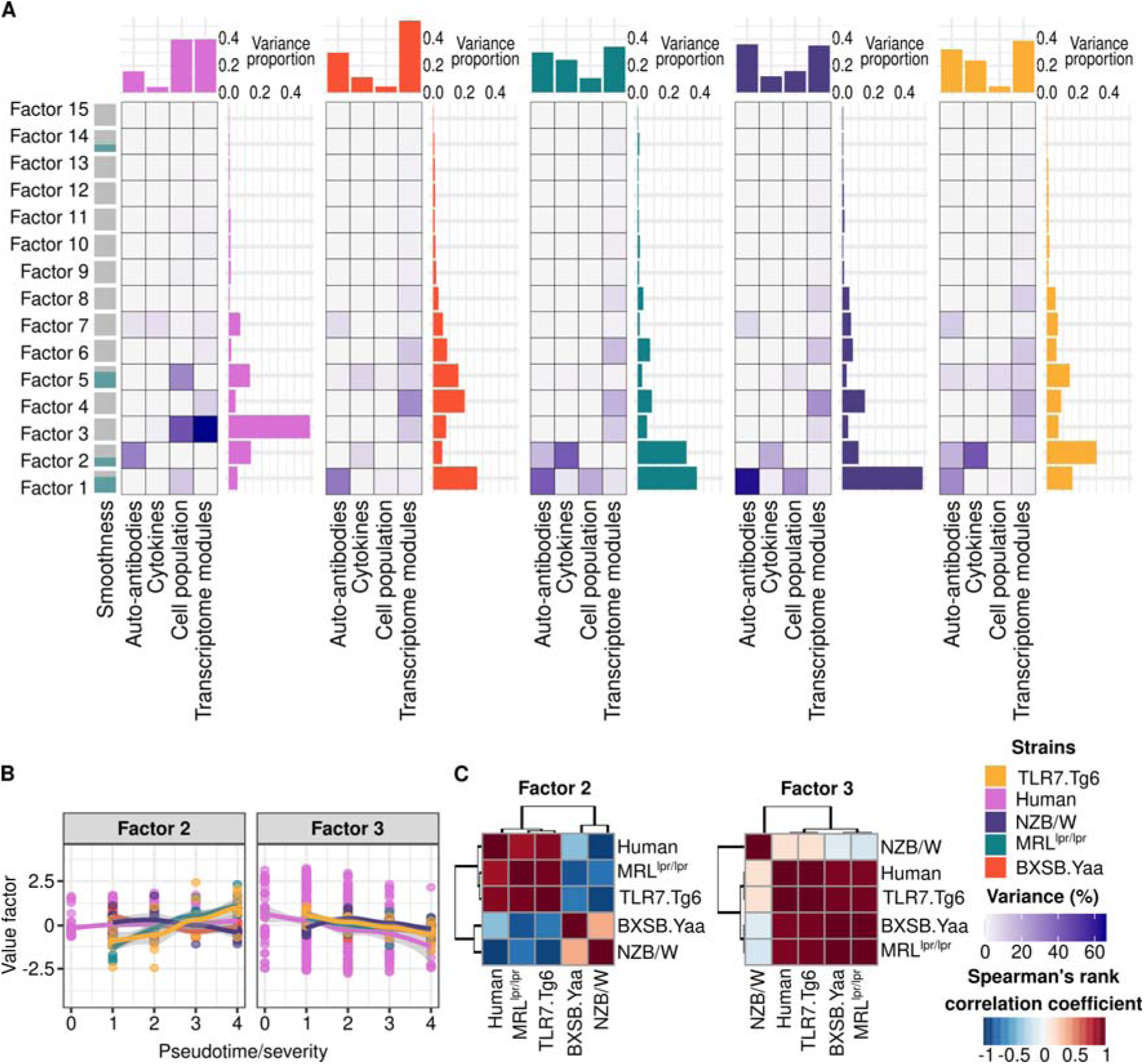
Longitudinal molecular factors are comparable between mouse models and SLE patients. **(A)** Explained variance proportion is shown per factor (rows), molecular layer of information (columns), human and mouse strain (panels). Factors’ longitudinal trends are represented on the left side of the representation in terms of smoothness. **(B)** Trend along severity is shown for each significant factor per group. Factor values per severity and group are summarized by means of loess regression. Trend along severity is shown for each significant factor per group. Factor values per severity and group are summarized by means of loess regression. **(C)** Factor value correlations between human and mouse models are depicted for the significant factors. Mouse models are color-coded as follows: yellow (Tlr7.Tg6), purple (NZB/W), green (MRL*^lpr/lpr^*), and red (BXSB.*Yaa*). Human results are colored pink.

Factor 1 increased over time in all mouse models (Fig. S8C), driven by the expansion of activated B cells (ABCs and IgM-IgD-B cells), memory T cells, neutrophils, and the depletion of naïve T cells (Fig. S9A). Cell proportion profiles in the spleen primarily originate from the MRL*^lpr/lpr^* and NZB/W mouse models (Fig. 3A), while the presence of anti-dsDNA, anti-RNP, and anti-histone autoantibodies was consistent across all mouse models within this factor (Fig. S9B). Factor 5 showed positive trends in the MRL*^lpr/lpr^*, BXSB.*Yaa*, and Tlr7.Tg6 models (Fig. S8B), distinguishing them from the NZB/W model and from humans (Fig. S8C). Cell populations and cytokines played major roles in this factor. Specifically, CD4+ T cells and CD4+/CD8+ double positive T cells (DP T cells) drove the factor in the Tlr7.Tg6 model (Fig. 3A and Fig. S9A), whereas myeloid-related cytokines (MCP-2, MIP-1α, IL-6, and TNF-α) predominantly influenced the factor in the BXSB.*Yaa* model (Fig. S9C).

Factors 2 and 3 were shared between humans and mice. Factor 2 was primarily influenced by autoantibodies against extractable nuclear antigens, particularly anti-SSA/Ro (Fig. S9B), which drove the positive trend correlations between humans, the MRL*^lpr/lpr^* and the Tlr7.Tg6 mouse models (Fig. 3B-C). Notably, these two mouse models contributed significantly to the explained variance (Fig. 3A) through the overexpression of cytokines measured in both species, such as IP-10, MIP-1β, IL-6, IL-17A and IL-22 (Fig. S9C). These cytokines have previously been associated with the regulation of interferon production, as well as other cytokines contributing to this factor that were only profiled in mice (Fig. S9C), including IFN-γ and IFN-λ (IL-28). Based on the SLEDAI decomposition into its clinical domains, it was found that this interferon-driven factor 2 was associated with constitutional and hematological clinical domains (Fig. 4A). Factor 3 showed significant downward trends in all the models correlating with humans (Fig. 3C), except NZB/W (Fig. 3B). Factor 3 was the factor explaining the most variance in humans (Fig. 3A), mainly driven by neutrophil expansion (Fig. S9A) and myeloid inflammatory and leukocyte activation-related transcriptional module upregulation (Fig. S9D). The cytokines contributing to the factor included MMP-8, TGFB1, IL-1Ra (Fig. S9C), which are known to be released by neutrophils upon stimulation, and BLC (*CXCL13* gene), which has been associated with nephritis in both humans and mice (36). In this line, the factor showed a strong association with the renal and cutaneous SLEDAI domains (Fig. 4A).

**Fig. 4:**
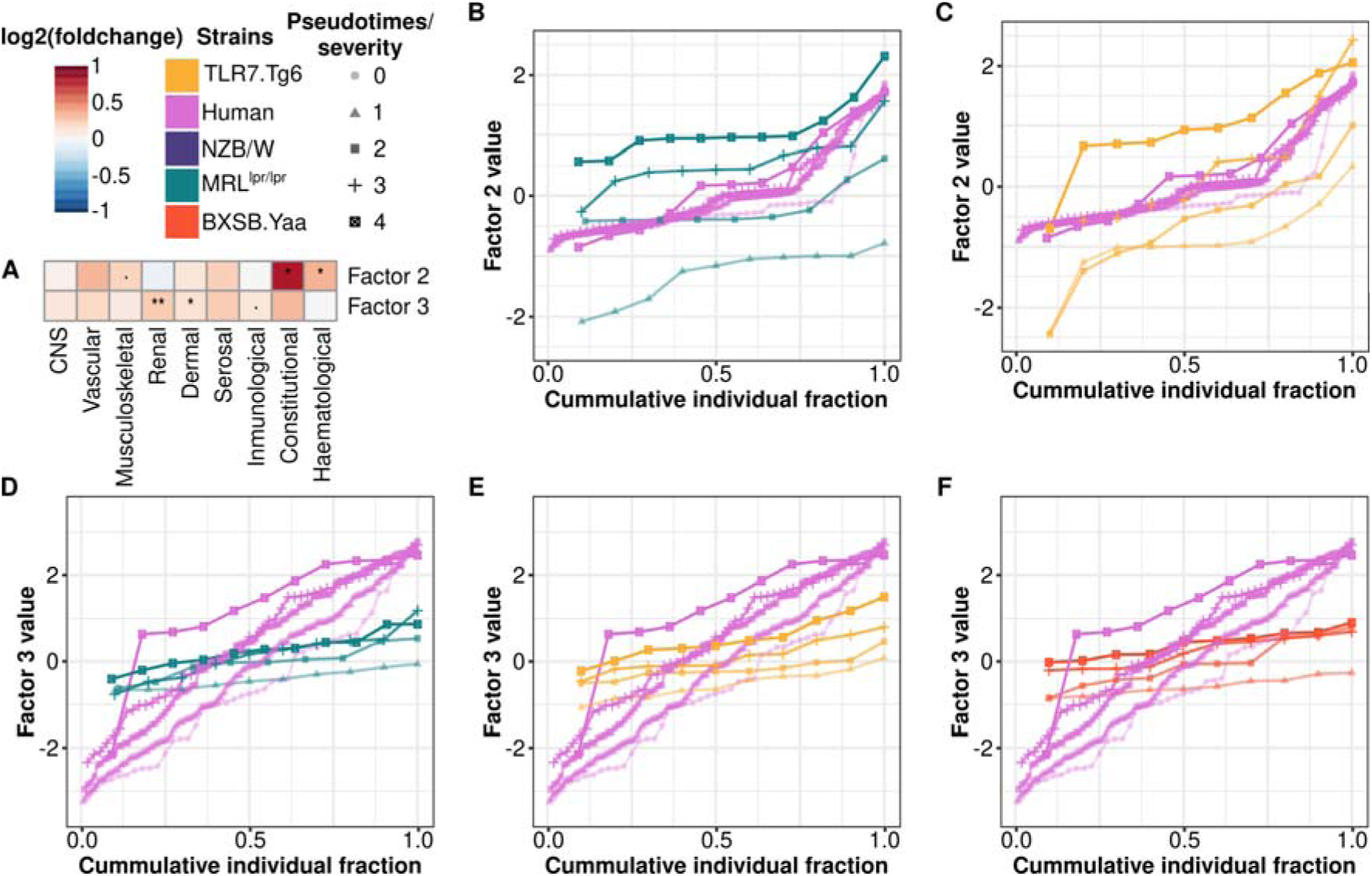
Pathogenic factors associate with specific clinical domains and time points in the different mouse models. **(A)** log_2_(fold change) between significant factor values and SLEDAI clinical domains values are shown (Wilcoxon signed rank test: .p-value < 0.1, *p-value < 0.05, **p-value < 0.01). The cumulative distributions per severity/pseudotime of individual fractions based on the factor values are depicted by mouse model and compared with human: **(B)** Factor 2 for MRL*^lpr/lpr^*, **(C)** Factor 2 for Tlr7.Tg6, **(D)** Factor 3 for, **(E)** Factor 3 for Tlr7.Tg6 and **(F)** Factor 3 for BXSB.*Yaa*. Mouse models are color-coded as follows: yellow (Tlr7.Tg6), purple (NZB/W), green (MRL*^lpr/lpr^*), and red (BXSB.*Yaa*). Human results are colored pink.

To identify the optimal time points for investigating human-related factors in mouse models, cumulative factor value distributions were plotted (Fig. 4B-F). For the interferon-related factor, MRL^lpr/lpr^ pseudotimes 2-3 (weeks 12-14) and Tlr7.Tg6 pseudotime 3 (week 16) closely resembled the distribution observed in severe SLE patients (Fig. 4B-C). For the neutrophil-mediated factor, none of the models matched the most severe human profiles at the sampled time points, indicating the need for longer time points to replicate the molecular dysregulation of this pathway in human SLE (Fig. 4D-F).

## DISCUSSION

Animal-based experimental setups are challenging due to heterogeneity, cost, time and ethical concerns. And poses the limitation that the outcomes may be unsatisfactory if models fail to molecularly mimic the intended disease. This likely influenced the FDA’s decision to allow human drug trials without prior preclinical animal testing. Nonetheless, animal models have been useful in understanding immune regulation, molecular disease mechanisms (37) and assessing the efficacy of drug candidates prior to human clinical trials (9). For complex disorders like SLE, the molecular dysregulations driving phenotypic similarities in animal models remain elusive. Therefore, a deeper understanding of these mechanisms in comparison with human disease is crucial for an efficient use of animal models. In this study, we longitudinally profiled different molecular layers in four spontaneous SLE mouse models and integrated with human data. The rationale behind this approach lies in the escalating molecular dysregulations observed with disease progression and potential molecular commonalities between mice and humans (Fig. 5). Thus, molecular information related with disease severity, defined as time in mice and activity in humans, was used in the integration analysis.

**Fig 5:**
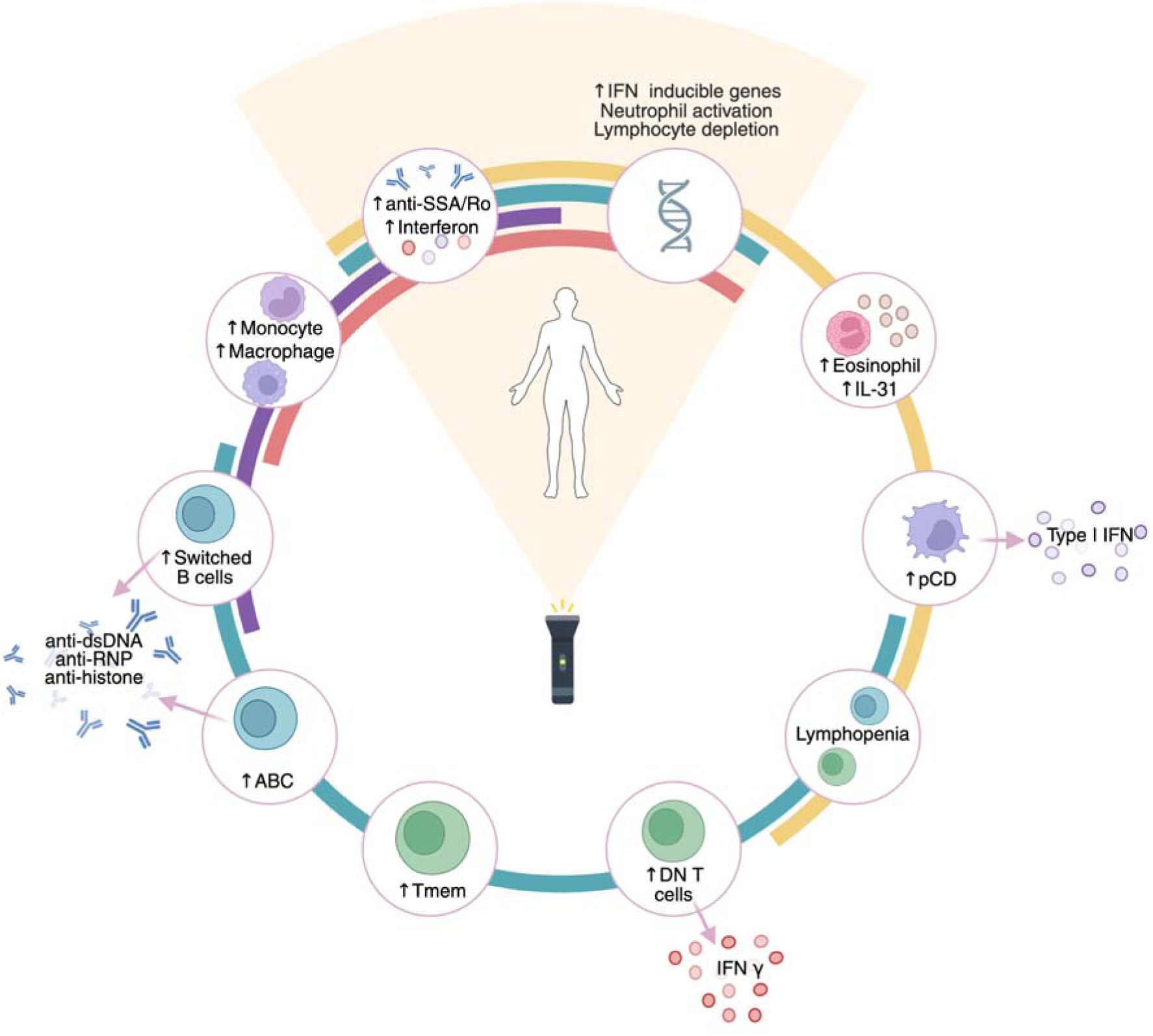
Molecular commonalities and specificities between SLE patients and spontaneous mouse models. Summary of the molecular signatures associated with the development of phenotypes in mouse models and the severity in human SLE. Molecular features associated with human SLE are depicted at 12 o’clock position in the plot. Each colored circular line represents a different mouse model: yellow (Tlr7.Tg6), purple (NZB/W), green (MRL*^lpr/lpr^*), and red (BXSB.*Yaa*). The surrounding molecular dysregulations were found to be associated with the development of the phenotype in at least one mouse model, defined by the circular lines.

Our results showed that the severity and 50% mortality rates of the mouse models were linked to the magnitude and timing of molecular dysregulations even after trying to align the samplings based on the severity of each model. The MRL*^lpr/lpr^* model, characterized by its aggressiveness with 50% mortality rate at 5 months in females (38), showed the highest number of differentially expressed molecular entities and early dysregulations, likely contributing to its rapid and severe phenotype. In contrast, the BXSB.*Yaa* and NZB/W models, with milder phenotypes, displayed fewer differentially expressed genes in the spleen and later kidney, correlating with 50% mortality rates between 5-8 months (38,39). The least aggressive Tlr7.Tg6 model, with 50% mortality in over 10 months (10), had the fewest differences, appearing only at the latest time point in the kidney. This supports the notion that disease severity is correlated with the extent and timing of molecular dysregulations. Moreover, specific temporal relationships between molecular layers and tissues were identified. Our findings indicated that molecular dysregulations initiate at the spleen level, followed by autoantibody production, and finally affect target tissues like the kidney. This sequential escalation of events is mirrored at the functional level, where, for example, dysregulated immune pathways precede apoptosis triggering in the tissues.

While spleen is often used in mouse immune studies, whole blood is utilized in human research due to accessibility as target tissue proxy. Our findings showed good transcriptional correlations between blood and spleen in mice, suggesting that blood is a reliable proxy for the spleen. However, neither fully captured the kidney’s molecular signature, as not all immune circulating cell types infiltrate tissues during the autoimmune process. Thus, it is reasonable to expect that not every immunological signature found in the spleen or blood would be present in the target organ.

Mouse model analyses uncovered various molecular dysregulations contributing to phenotype development showing continuous trends over time. These molecular dysregulations were consolidated within factor 1 during integration analysis. This factor primarily involved immune cell migration and activation across different cell compartments. The B cell compartment showed an activation process, including an expansion of switched IgM-IgD-B cells (mainly in MRL^lpr/lpr^ and NZB/W models) and ABCs (exclusive to MRL^lpr/lpr^) in the spleen. These cell types may be responsible for the overproduction of anti-dsDNA, anti-RNP, and anti-histone autoantibodies also associated with this factor. Given the emerging significance of ABCs in SLE, our findings suggest that the MRL*^lpr/lpr^*model is particularly suited for studying the role of ABCs expansion in SLE pathogenesis (40). In addition, CD4+ T cell lymphopenia was identified, mirroring the characteristic reduction in CD4+ T cell compartments observed in SLE patients (41), marked by a decrease in naïve T cell populations and an increase in memory T cells. This process was particularly noted in the MRL*^lpr/lpr^* model and in the Tlr7.Tg6 at later time points. Unfortunately, these molecular processes identified in mice were not fully recapitulated by the available human molecular data, leaving their relevance in human disease unclear. Nevertheless, one of the factors shared between mice and humans (factor 3) included the downregulation in CD4+ and CD8+ T cell compartments, which might be associated with a reduction in their naïve compartments and related with the lymphopenia.

Certain molecular signatures identified during regression analyses were not encompassed by any factor during integration. For instance, DN T cells were found to be dysregulated very early in the MRL*^lpr/lpr^*model, but they did not align with any factor associated with disease progression. We posit that this is due to a sustained and significant expansion of these cell types, thereby not showing a longitudinal trend. However, this may indicate their crucial role during the initiation of disease in the mice. The MRL*^lpr/lpr^* model is recognized for presenting impaired apoptosis via the Fas-FasL pathway (42), leading to the escape from activation-induced cell death of these DN T cells. Also, the expansion of memory T cells in this model may be attributed to impaired T cell apoptosis. Both processes have previously been related to SLE pathogenesis (27,43). A similar scenario was observed for pDCs in the Tlr7.Tg6 model, where their expansion and overactivation are linked to the upregulation of the *Tlr7* gene (44), yet pDCs were not associated with any factor in our study. Notably, this model showed an expansion of eosinophils and the overexpression of IL-31 and interferon-related proteins, which have been linked to inflammatory skin diseases (45,46). Thus, the Tlr7.Tg6 model may serve as a promising candidate for studying cutaneous manifestations of lupus and their treatment; however, this particular profile was not directly associated with disease severity in neither humans nor mouse models, at least not with severity as defined in this analysis. In the case of NZB/W and BXSB.*Yaa*, monocyte/macrophage-related signatures may be associated with the phenotype development in the former, due to an increased proportion of monocytes in the spleen, and based on cytokine and transcriptomic signatures in the latter. However, no factor reflected this association or its sharedness with human molecular signatures. Notably, although monocytosis is a characteristic feature of the BXSB.*Yaa* model, it is not associated with disease severity, unlike in the NZB/W model (47). This may explain the lack of association observed in our severity-oriented analysis.

Whole-blood human transcriptome analyses have consistently revealed dysregulations in various molecular signatures associated with SLE. These include upregulation of interferon-inducible genes, activation of neutrophils, and lymphocyte depletion in both T and B cell compartments (4,48–50). The cross-species integration revealed, for the first time, a core of shared molecular features representing these well-known SLE-associated signatures between species. The interferon signature stands out as one of the most studied immune pathways linked to human SLE. Cytokine profiling, the established relationship between interferon production and anti-SSA/Ro autoantibodies (51,52), and the absence of cell proportions contributing to this factor led us to deduce that factor 2 reflects this interferon-inducible signature in both humans and mice. Moreover, the minimal differences observed in its cumulative distributions across human severities align with previous findings regarding the persistence of the interferon signature even after disease remission (50). Based on our results, both the MRL*^lpr/lpr^*and Tlr7.Tg6 models are suitable for studying phenotypes driven by interferon dysregulation. The time points utilized in our study align well with the range required to examine both late and early molecular changes leading to this interferon signature. Furthermore, our study unveiled an interferon dysregulation cascade in the MRL*^lpr/lpr^* model, initiated by the overproduction of IFN-γ possibly stemming from the expansion of DN T cells in the model, which have been linked to kidney damage in SLE (27). Conversely, in the Tlr7.Tg6 model, interferon production likely arises from expanded pDCs, known as major producers of type I interferons, and previously associated with disease activity in cutaneous lupus (52). These findings suggest distinct initiation pathways for the interferon signature in these mouse models, each associated with different SLE phenotypes. However, further experiments are needed to elucidate whether both pathways coexist or trigger the interferon signature in different subgroups of SLE patients.

Another significant peripheral blood molecular signature commonly associated with SLE patients involves neutrophil activation and/or expansion, often linked to the proliferation of a pathogenic subset of activated neutrophils known as low-density granulocytes (53). This signature has been consistently linked to lupus nephritis (4–6), underscoring its pathological relevance in the disease, and evidenced here by the factor-association with the renal SLEDAI domain. Factor 3 captured this molecular pathway through a pronounced expansion of neutrophils coupled with an overexpression in neutrophil-related cytokines and transcriptional modules. Additionally, another well-established SLE-related signature, marked by a general depletion of B and T lymphocytes, was also encompassed within this factor and reflected in the transcriptome. Unfortunately, although the factor was present in the BXSB.*Yaa*, Tlr7.Tg6, and MRL*^lpr/lpr^* models, none of them exhibited values reaching the highest levels observed in humans. Among the profiled time points, the sharpest upward trend was observed in the Tlr7.Tg6 model, suggesting it may be the most suitable model for profiling at longer time points, especially given its lower aggressiveness. Notably, recent findings have demonstrated extensive sharing of neutrophil transcriptional programs between mice and humans (54), further underpinning these results.

## CONCLUSIONS

Among the four mouse models analyzed, Tlr7.Tg6 and MRL*^lpr/lpr^*most closely mirror the interferon and neutrophil-related molecular dysregulations observed in human SLE. These models exhibit distinct molecular characteristics during pathogenesis and varying degrees of severity, thus offering utility depending on the specific experimental objectives or pathways under investigation. While BXSB.*Yaa* and NZB/W also demonstrated interesting signatures, such as monocyte/macrophage migration and activation or an early and heightened autoantibody production, none directly mirrored the molecular dysregulations observed in human SLE, at least based on our results. This study, for the first time, directly integrates molecular information from SLE mouse models within the context of human SLE, shedding light on which mouse models and time points most faithfully recapitulate the molecular signatures associated with the human disease. We anticipate that these results will help to delineate future SLE mouse model-based experimental designs, thereby enhancing the quality and relevance of their outcomes.

## ABBREVIATIONS

ABCs: Age-associated B cells
ACR: Albumin-to-creatinine ratio
ANAs: Antinuclear antibodies
anti-CENP-B: Anti-centromere protein B
anti-dsDNA: Anti-double-stranded DNA
anti-RNP: Anti-ribonucleoprotein
anti-Sm: Anti-Smith antigen
anti-SSA/La: Anti-Sjögren’s-syndrome-related antigen A
anti-SSB/Ro: Anti-Sjögren’s-syndrome-related antigen B
CXCL-1: Chemokine (C-X-C motif) ligand 1
DN T cells: Double-negative T cells
DP T cells: Double-positive T cells
EDTA: Ethylenediaminetetraacetic acid
ENA-78: Epithelial-derived neutrophil-activating peptide 78
FDA: Food and Drug Administration
FDR: False discovery rate
G-CSF: Granulocyte colony-stimulating factor
GM-CSF: Granulocyte-macrophage colony-stimulating factor
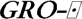: Growth-related oncogene-alpha
GSEA: Gene Set Enrichment Analysis
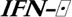: Interferon-alpha
IFN-J: Interferon-gamma
IgG: Immunoglobulin G
IgM-IgD-B cells: Class-switched B cells
IL: Interleukin
IP-10: Interferon gamma-induced protein 10
MCP-1: Monocyte chemoattractant protein-1
MCP-3: Monocyte chemoattractant protein-3
M-CSF: Monocyte-colony stimulating factor
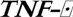: Macrophage inflammatory protein-1 alpha
MIP-1ß: Macrophage inflammatory protein-1 beta
MIP-2: Macrophage inflammatory protein-2
MRL: Murphy Roths Large
MZ B cells: Marginal zone B cells
NZB: New Zealand Black
NZW: New Zealand White
PBS: Phosphate-buffered saline
PCs: Plasma cells
pDCs: Plasmacytoid dendritic cells
RANTES: Regulated upon activation, normal T cell expressed and secreted
SLE: Systemic lupus erythematosus
T3 B cells: Transitional T3 B cells
Tmem: Memory T cells
TN: Naïve T cells
[inline]: Tumor necrosis factor-alpha

## DECLARATIONS

### Ethics approval and consent to participate

The Ethical Review Boards of the 18 participating institutions approved the protocol of the study: Referral Center for Systemic Autoimmune Diseases, Fondazione IRCCS Ca’ Granda Ospedale Maggiore Policlinico di Milano (IRCCS); Centre Hospitalier Universitaire de Brest, Hospital de la Cavale Blanche (UBO); Pôle de pathologies rhumatismales systémiques et inflammatoires, Institut de Recherche Expérimentale et Clinique Université catholique de Louvain (UCL); Centro Hospitalar do Porto (CHP); Servicio Cantabro de Salud, Hospital Universitario Marqués de Valdecilla (SCS); Hospital Clinic I Provicia, Institut d’Investigacions Biomèdiques August Pi i Sunyer (IDIBAPS); Katholieke Universiteit Leuven (KU.LEUVEN); Klinikum der Universitaet zu Koeln, Cologne (UKK); Medizinische Hochschule Hannover (MHH); Medical University Vienna (MUW); Servicio Andaluz de Salud, Hospital Universitario Reina Sofía (SAS.Co); Servicio Andaluz de Salud, Complejo hospitalario Universitario de Granada, Hospital Universitario San Cecilio (SAS.GrE); Servicio Andaluz de Salud, Complejo hospitalario Universitario de Granada, Hospital Virgen de las Nieves (SAS.GrN); Servicio Andaluz de Salud, Hospital Regional Universitario de Málaga (SAS.Ma); Università degli studi di Milano (UNIMI); Hospitaux Universitaires de Genève (UNIGE); University of Szeged (USC); Charité (DRFZ). The study adhered to the standards set by the International Conference on Harmonization and Good Clinical Practice (ICH-GCP), and to the ethical principles that have their origin in the Declaration of Helsinki. The protection of the confidentiality of records that could identify the included subjects is ensured as defined by the EU Directive 2001/20/EC and the applicable national and international requirements relating to data protection in each participating country. All patients signed the informed consent prior to recruitment. Both centers involved in the mouse model sampling (FPS and DRFZ) obtained an ethical or animal use approval from their local ethical or experimental animal committee, and for Spain, the Ministry of Agriculture.

### Consent for publication

Not applicable

### Availability of data and materials

Human and mouse datasets are hosted by ELIXIR Luxembourg and will be available after publication. No custom code or unpublished methods were used in the study. The scripts used in the generation of this manuscript are available upon request.

### Author contributions

All authors revised the manuscript critically for important intellectual content, and approved the final version to be published. Design and conceptualization: MAR, FMD, GB, MM, ZM. Sample acquisition: MM, ZM, AB, LB, PCC. Data profiling: ZM, AB, CM, MOB, EB, PFCC, JOP, RL. Methodology and statistical analysis: GB, MRT, JK, JL, IP, LB. Funding acquisition: MAR. Supervision: MAR, GB. Original draft: GB, MRT, MAR.

## Acknowledgements

The authors would like to particularly express their gratitude to the investigators of the PRECISESADS Clinical and Flow-Cytometry consortiums listed below, and to the patients, nurses and many others who helped directly or indirectly in the consecution of this study. This work was supported by ELIXIR Luxembourg.

PRECISESADS Clinical Consortium: Lorenzo Beretta, Barbara Vigone, Jacques-Olivier Pers, Alain Saraux, Valérie Devauchelle-Pensec, Divi Cornec, Sandrine Jousse-Joulin, Bernard Lauwerys, Julie Ducreux, Anne-Lise Maudoux, Carlos Vasconcelos, Ana Tavares, Esmeralda Neves, Raquel Faria, Mariana Brandão, Ana Campar, António Marinho, Fátima Farinha, Isabel Almeida, Miguel Angel Gonzalez-Gay, Ricardo Blanco Alonso, Alfonso Corrales Martínez, Ricard Cervera, Ignasi Rodríguez-Pintó, Gerard Espinosa, Rik Lories, Ellen De Langhe, Nicolas Hunzelmann, Doreen Belz, Torsten Witte, Niklas Baerlecken,

Georg Stummvoll, Michael Zauner, Michaela Lehner, Eduardo Collantes, Rafaela Ortega-Castro, Mª Angeles Aguirre-Zamorano, Alejandro Escudero-Contreras, Mª Carmen Castro-Villegas, Yolanda Jiménez Gómez, Norberto Ortego, María Concepción Fernández Roldán, Enrique Raya, Inmaculada Jiménez Moleón, Enrique de Ramon, Isabel Díaz Quintero, Pier Luigi Meroni, Maria Gerosa, Tommaso Schioppo, Carolina Artusi, Carlo Chizzolini, Aleksandra Dufour, Donatienne Wynar, Laszló Kovács, Attila Balog, Magdolna Deák, Márta Bocskai, Sonja Dulic, Gabriella Kádár, Falk Hiepe, Velia Gerl, Silvia Thiel, Manuel Rodriguez Maresca, Antonio López-Berrio, Rocío Aguilar-Quesada, Héctor Navarro-Linares, Yiannis Ioannou, Chris Chamberlain, Jacqueline Marovac.

PRECISESADS Flow-Cytometry Consortium: Christophe Jamin, Concepción Marañón, Lucas Le Lann, Quentin Simon, Bénédicte Rouvière, Nieves Varela, Brian Muchmore, Aleksandra Dufour, Montserrat Alvarez,Carlo Chizzolini, Jonathan Cremer, Ellen De Langhe, Nuria Barbarroja, Chary Lopez-Pedrera, Velia Gerl, Laleh Khodadadi, Qingyu Cheng, Anne Buttgereit, Zuzanna Makowska, Aurélie De Groof, Julie Ducreux, Elena Trombetta, Tianlu Li, Damiana Alvarez-Errico, Torsten Witte, Katja Kniesch, Nancy Azevedo, Esmeralda Neves, Sambasiva Rao, Pierre-Emmanuel Jouve, Jacques-Olivier Pers.

## ADDITIONAL FILES

Additional File 1 (xlsx): Statistical analyses for MRL^lpr/lpr^. Statistics from regression and correlation analyses are included in the file.

Additional File 2 (xlsx): Statistical analyses for BXSB.*Yaa*. Statistics from regression and correlation analyses are included in the file.

Additional File 3 (xlsx): Statistical analyses for NZB/W. Statistics from regression and correlation analyses are included in the file.

Additional File 4 (xlsx): Statistical analyses for Tlr7.Tg6. Statistics from regression and correlation analyses are included in the file.

Additional File 5 (xlsx): Statistical analyses for human samples. Statistics from regression and correlation analyses are included in the file.

## REFERENCES

1. Wadman M. FDA no longer has to require animal testing for new drugs. Science. 2023 Jan 13;379(6628):127–8.

2. Barturen G, Beretta L, Cervera R, Van Vollenhoven R, Alarcón-Riquelme ME. Moving towards a molecular taxonomy of autoimmune rheumatic diseases. Nat Rev Rheumatol. 2018 Jan 24;14(2):75–93.

3. Kaul A, Gordon C, Crow MK, Touma Z, Urowitz MB, van Vollenhoven R, et al. Systemic lupus erythematosus. Nat Rev Dis Primer. 2016 Jun 16;2(1):1–21.

4. Banchereau R, Hong S, Cantarel B, Baldwin N, Baisch J, Edens M, et al. Personalized Immunomonitoring Uncovers Molecular Networks that Stratify Lupus Patients. Cell. 2016 Apr 21;165(3):551–65.

5. Toro-Domínguez D, Martorell-Marugán J, Goldman D, Petri M, Carmona-Sáez P, Alarcón-Riquelme ME. Stratification of Systemic Lupus Erythematosus Patients Into Three Groups of Disease Activity Progression According to Longitudinal Gene Expression. Arthritis Rheumatol. 2018;70(12):2025–35.

6. Barturen G, Babaei S, Català-Moll F, Martínez-Bueno M, Makowska Z, Martorell-Marugán J, et al. Integrative Analysis Reveals a Molecular Stratification of Systemic Autoimmune Diseases. Arthritis Rheumatol. 2021;73(6):1073–85.

7. Mahieu MA, Strand V, Simon LS, Lipsky PE, Ramsey-Goldman R. A critical review of clinical trials in systemic lupus erythematosus. Lupus. 2016 Sep 1;25(10):1122–40.

8. Perry D, Sang A, Yin Y, Zheng YY, Morel L. Murine Models of Systemic Lupus Erythematosus. BioMed Res Int. 2011 Feb 14;2011:e271694.

9. Moore E, Putterman C. Are lupus animal models useful for understanding and developing new therapies for human SLE? J Autoimmun. 2020 Aug 1;112:102490.

10. Deane JA, Pisitkun P, Barrett RS, Feigenbaum L, Town T, Ward JM, et al. Control of Toll-like Receptor 7 Expression Is Essential to Restrict Autoimmunity and Dendritic Cell Proliferation. Immunity. 2007 Nov 26;27(5):801–10.

11. Li W, Titov AA, Morel L. An update on lupus animal models. Curr Opin Rheumatol. 2017 Sep;29(5):434.

12. Martin M. Cutadapt removes adapter sequences from high-throughput sequencing reads. EMBnet.journal. 2011 May 2;17(1):10–2.

13. Dobin A, Davis CA, Schlesinger F, Drenkow J, Zaleski C, Jha S, et al. STAR: ultrafast universal RNA-seq aligner. Bioinformatics. 2013 Jan 1;29(1):15–21.

14. Li B, Dewey CN. RSEM: accurate transcript quantification from RNA-Seq data with or without a reference genome. BMC Bioinformatics. 2011 Aug 4;12(1):323.

15. Frankish A, Diekhans M, Ferreira AM, Johnson R, Jungreis I, Loveland J, et al. GENCODE reference annotation for the human and mouse genomes. Nucleic Acids Res. 2019 Jan 8;47(D1):D766–73.

16. Jamin C, Le Lann L, Alvarez-Errico D, Barbarroja N, Cantaert T, Ducreux J, et al. Multi-center harmonization of flow cytometers in the context of the European “PRECISESADS” project. Autoimmun Rev. 2016 Nov 1;15(11):1038–45.

17. Becton, Dickinson and Company. FlowJo^TM^ Software. Version 107. 2023;

18. Love MI, Huber W, Anders S. Moderated estimation of fold change and dispersion for RNA-seq data with DESeq2. Genome Biol. 2014 Dec 5;15(12):550.

19. Hänzelmann S, Castelo R, Guinney J. GSVA: Gene set variation analysis for microarray and RNA-Seq data. BMC Bioinformatics. 2013;14:undefined-undefined.

20. Rinchai D, Roelands J, Toufiq M, Hendrickx W, Altman MC, Bedognetti D, et al. BloodGen3Module: blood transcriptional module repertoire analysis and visualization using R. Bioinformatics. 2021 Feb 24;37(16):2382–9.

21. Sayers EW, Bolton EE, Brister JR, Canese K, Chan J, Comeau DC, et al. Database resources of the national center for biotechnology information. Nucleic Acids Res. 2022 Jan 7;50(D1):D20–6.

22. Wu T, Hu E, Xu S, Chen M, Guo P, Dai Z, et al. clusterProfiler 4.0: A universal enrichment tool for interpreting omics data. Innov Camb Mass. 2021 Aug 28;2(3):100141.

23. Liberzon A, Birger C, Thorvaldsdóttir H, Ghandi M, Mesirov JP, Tamayo P. The Molecular Signatures Database Hallmark Gene Set Collection. Cell Syst. 2015 Dec 23;1(6):417–25.

24. Velten B, Braunger JM, Argelaguet R, Arnol D, Wirbel J, Bredikhin D, et al. Identifying temporal and spatial patterns of variation from multimodal data using MEFISTO. Nat Methods. 2022 Feb;19(2):179–86.

25. Mosca M, Bombardieri S. Assessing remission in systemic lupus erythematosus. Clin Exp Rheumatol. 2006;24(6 Suppl 43):S-99–104.

26. Petri M, Genovese M, Engle E, Hochberg M. Definition, incidence, and clinical description of flare in systemic lupus erythematosus. A prospective cohort study. Arthritis Rheum. 1991;34(8):937–44.

27. Crispín JC, Oukka M, Bayliss G, Cohen RA, Van Beek CA, Stillman IE, et al. Expanded Double Negative T Cells in Patients with Systemic Lupus Erythematosus Produce IL-17 and Infiltrate the Kidneys1. J Immunol. 2008 Dec 15;181(12):8761–6.

28. Cong M, Liu T, Tian D, Guo H, Wang P, Liu K, et al. Interleukin-2 Enhances the Regulatory Functions of CD4(+)T Cell-Derived CD4(-)CD8(-) Double Negative T Cells. J Interferon Cytokine Res Off J Int Soc Interferon Cytokine Res. 2016 Aug;36(8):499–505.

29. Fahey TJ, Tracey KJ, Tekamp-Olson P, Cousens LS, Jones WG, Shires GT, et al. Macrophage inflammatory protein 1 modulates macrophage function. J Immunol Baltim Md 1950. 1992 May 1;148(9):2764–9.

30. Rot A, Krieger M, Brunner T, Bischoff SC, Schall TJ, Dahinden CA. RANTES and macrophage inflammatory protein 1 alpha induce the migration and activation of normal human eosinophil granulocytes. J Exp Med. 1992 Dec 1;176(6):1489–95.

31. Horejs-Hoeck J, Schwarz H, Lamprecht S, Maier E, Hainzl S, Schmittner M, et al. Dendritic cells activated by IFN-γ/STAT1 express IL-31 receptor and release proinflammatory mediators upon IL-31 treatment. J Immunol Baltim Md 1950. 2012 Jun 1;188(11):5319–26.

32. Cheung PFY, Wong CK, Ho AWY, Hu S, Chen DP, Lam CWK. Activation of human eosinophils and epidermal keratinocytes by Th2 cytokine IL-31: implication for the immunopathogenesis of atopic dermatitis. Int Immunol. 2010 Jun;22(6):453–67.

33. Jia T, Serbina NV, Brandl K, Zhong MX, Leiner IM, Charo IF, et al. Additive Roles for MCP-1 and MCP-3 in CCR2-mediated Recruitment of Inflammatory Monocytes During Listeria monocytogenes Infection. J Immunol Baltim Md 1950. 2008 May 15;180(10):6846–53.

34. Chen S, Saeed AFUH, Liu Q, Jiang Q, Xu H, Xiao GG, et al. Macrophages in immunoregulation and therapeutics. Signal Transduct Target Ther. 2023 May 22;8(1):1–35.

35. Pajulas A, Zhang J, Kaplan MH. The World according to IL-9. J Immunol. 2023 Jul 1;211(1):7– 14.

36. Worthmann K, Gueler F, von Vietinghoff S, Davalos-Mißlitz A, Wiehler F, Davidson A, et al. Pathogenetic role of glomerular CXCL13 expression in lupus nephritis. Clin Exp Immunol. 2014 Oct;178(1):20–7.

37. Reza Khorramizadeh M, Saadat F. Animal models for human disease. Anim Biotechnol. 2020;153–71.

38. Andrews BS, Eisenberg RA, Theofilopoulos AN, Izui S, Wilson CB, McConahey PJ, et al. Spontaneous murine lupus-like syndromes. Clinical and immunopathological manifestations in several strains. J Exp Med. 1978 Nov 1;148(5):1198–215.

39. Ibnou-Zekri N, Iwamoto M, Fossati L, McConahey PJ, Izui S. Role of the major histocompatibility complex class II Ea gene in lupus susceptibility inlmice. Proc Natl Acad Sci. 1997 Dec 23;94(26):14654–9.

40. Mouat IC, Goldberg E, Horwitz MS. Age-associated B cells in autoimmune diseases. Cell Mol Life Sci. 2022 Jul 7;79(8):402.

41. Yuan S, Zeng Y, Li J, Wang C, Li W, He Z, et al. Phenotypical changes and clinical significance of CD4+/CD8+ T cells in SLE. Lupus Sci Med. 2022 Jun 1;9(1):e000660.

42. Singer GG, Carrera AC, Marshak-Rothstein A, Martínez-A C, Abbas AK. Apoptosis, fas and systemic autoimmunity: the MRL-*Ipr/Ipr* model. Curr Opin Immunol. 1994 Dec 1;6(6):913–20.

43. Suárez-Fueyo A, Barber DF, Martínez-Ara J, Zea-Mendoza AC, Carrera AC. Enhanced Phosphoinositide 3-Kinase δ Activity Is a Frequent Event in Systemic Lupus Erythematosus That Confers Resistance to Activation-Induced T Cell Death. J Immunol. 2011 Sep 1;187(5):2376–85.

44. Di Domizio J, Blum A, Gallagher-Gambarelli M, Molens JP, Chaperot L, Plumas J. TLR7 stimulation in human plasmacytoid dendritic cells leads to the induction of early IFN-inducible genes in the absence of type I IFN. Blood. 2009 Aug 27;114(9):1794–802.

45. Kuzumi A, Yoshizaki A, Matsuda KM, Kotani H, Norimatsu Y, Fukayama M, et al. Interleukin-31 promotes fibrosis and T helper 2 polarization in systemic sclerosis. Nat Commun. 2021 Oct 12;12(1):5947.

46. Guiducci C, Tripodo C, Gong M, Sangaletti S, Colombo MP, Coffman RL, et al. Autoimmune skin inflammation is dependent on plasmacytoid dendritic cell activation by nucleic acids via TLR7 and TLR9. J Exp Med. 2010 Nov 29;207(13):2931–42.

47. Rogers NJ, Gabriel L, Nunes CT, Rose SJ, Thiruudaian V, Boyle J, et al. Monocytosis in BXSB mice is due to epistasis between Yaa and the telomeric region of Chromosome 1 but does not drive the disease process. Genes Immun. 2007 Dec;8(8):619–27.

48. Bennett L, Palucka AK, Arce E, Cantrell V, Borvak J, Banchereau J, et al. Interferon and Granulopoiesis Signatures in Systemic Lupus Erythematosus Blood. J Exp Med. 2003 Mar 17;197(6):711–23.

49. Chaussabel D, Quinn C, Shen J, Patel P, Glaser C, Baldwin N, et al. A Modular Analysis Framework for Blood Genomics Studies: Application to Systemic Lupus Erythematosus. Immunity. 2008 Jul 18;29(1):150–64.

50. Toro-Domínguez D, Martorell-Marugán J, Martinez-Bueno M, López-Domínguez R, Carnero-Montoro E, Barturen G, et al. Scoring personalized molecular portraits identify Systemic Lupus Erythematosus subtypes and predict individualized drug responses, symptomatology and disease progression. Brief Bioinform. 2022 Sep 1;23(5):bbac332.

51. Amezcua-Guerra LM, Márquez-Velasco R, Chávez-Rueda AK, Castillo-Martínez D, Massó F, Páez A, et al. Type III Interferons in Systemic Lupus Erythematosus: Association Between Interferon λ3, Disease Activity, and Anti-Ro/SSA Antibodies. JCR J Clin Rheumatol. 2017 Oct;23(7):368.

52. Imgenberg-Kreuz J, Almlöf JC, Leonard D, Sjöwall C, Syvänen AC, Rönnblom L, et al. Shared and Unique Patterns of DNA Methylation in Systemic Lupus Erythematosus and Primary Sjögren’s Syndrome. Front Immunol [Internet]. 2019 Jul 30 [cited 2024 Mar 28];10. Available from: https://www.frontiersin.org/journals/immunology/articles/10.3389/fimmu.2019.01686/full

53. Mistry P, Nakabo S, O’Neil L, Goel RR, Jiang K, Carmona-Rivera C, et al. Transcriptomic, epigenetic, and functional analyses implicate neutrophil diversity in the pathogenesis of systemic lupus erythematosus. Proc Natl Acad Sci. 2019 Dec 10;116(50):25222–8.

54. Hackert NS, Radtke FA, Exner T, Lorenz HM, Müller-Tidow C, Nigrovic PA, et al. Human and mouse neutrophils share core transcriptional programs in both homeostatic and inflamed contexts. Nat Commun. 2023 Dec 8;14(1):8133.

