## Supplementary Materials for "Assessing the Molecular Validity of Spontaneous Lupus Mouse Models and Its Implication for Human Studies"

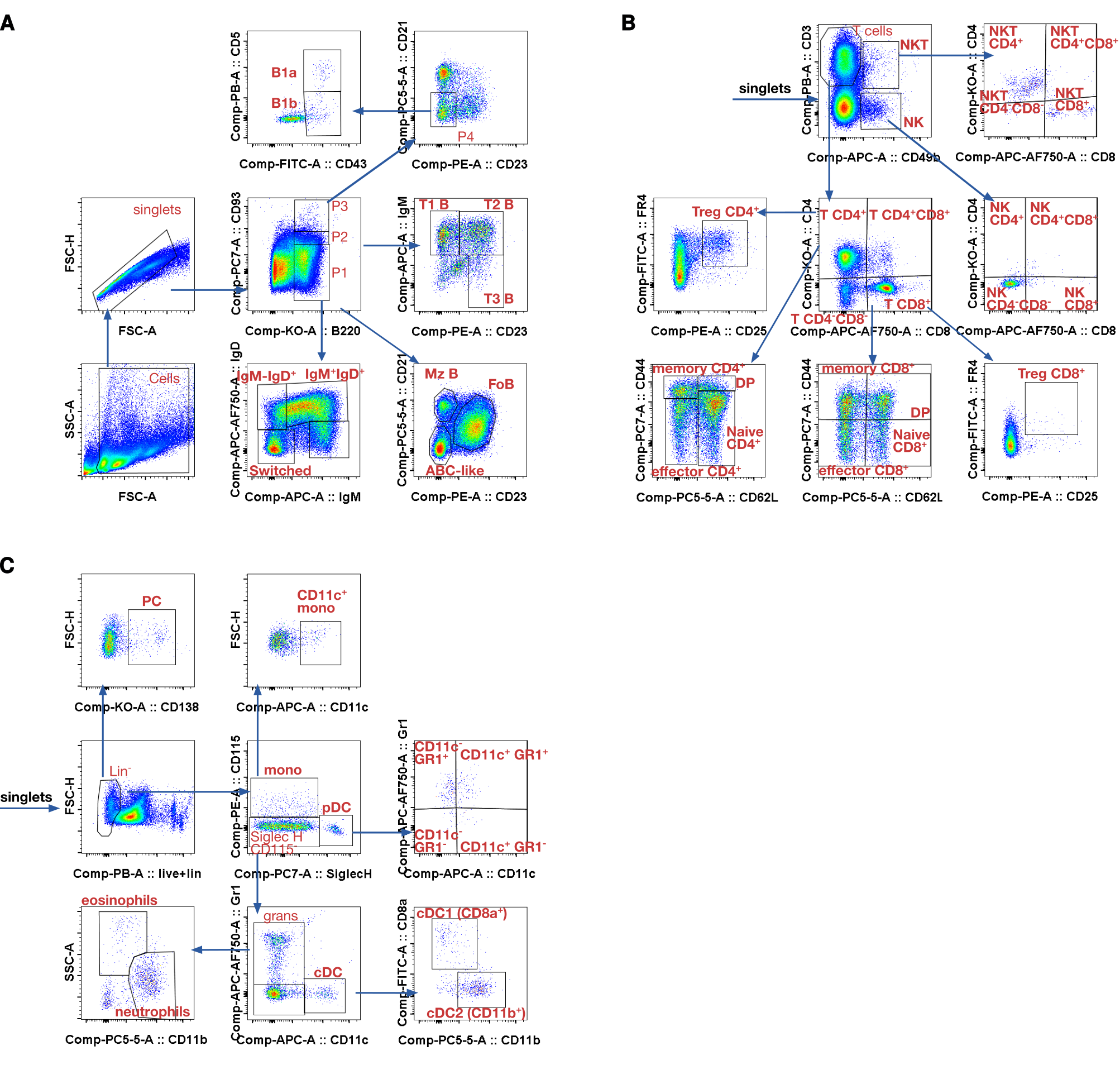
**Fig. S1: Mouse spleen cell population panels and gating.**

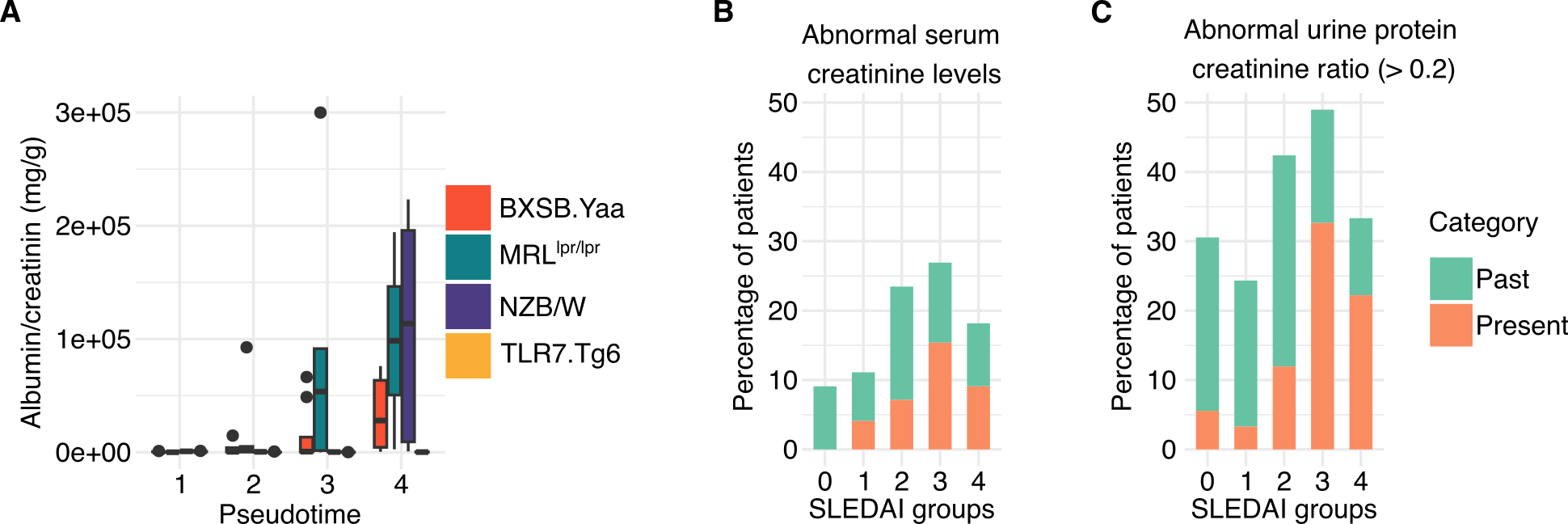
**Fig. S2: Markers of kidney involvement in lupus mouse models and human SLE patients.** (A) Urine albumin-to-creatinine ratio (ACR) measurements over pseudotimes (1,2,3 and 4) and mouse models: red (BXSB.*Yaa*), green (MRL*^lpr/lpr^*), purple (NZB/W) and yellow (Tlr7.Tg6). (B) Percentage of patients with abnormal serum creatinine levels, defined as serum creatinine levels ≥ 20% upper laboratory normal, serum creatinine levels ≥ 50% as compared with normal creatinine levels or reduced glomerular filtration rate (< 60 mL/min). (C) Percentage of patients with abnormal urine protein-to-creatinine ratio (> 0.2 mg/mg). Abnormalities were classified into present (within 4 weeks and not resolved) and past (observed and resolved or not verified since an observation of more than 4 weeks), and patients stratified by SLEDAI groups.

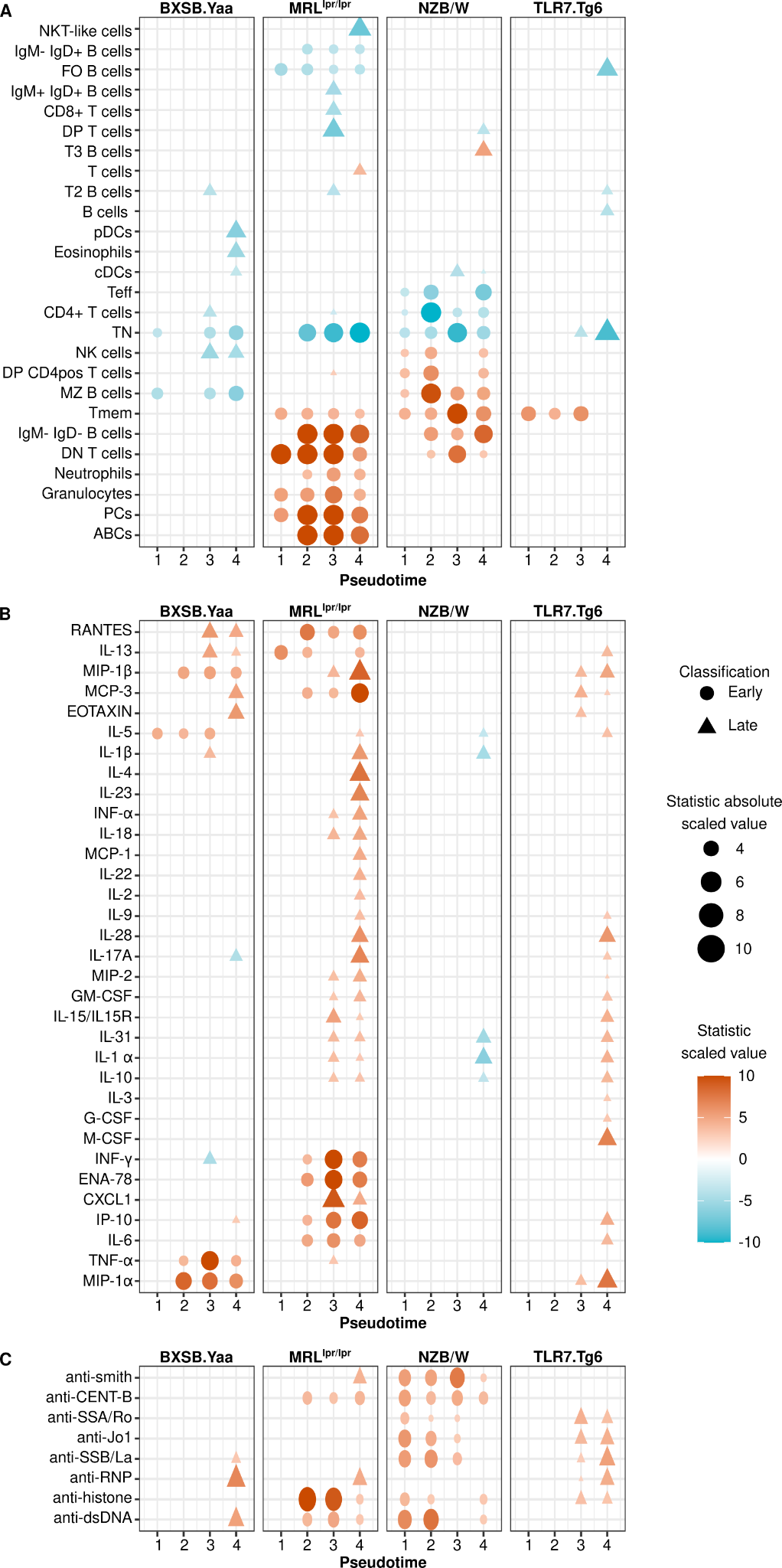

**Fig. S3: Serological time point analyses per mouse model.**

Cross-sectional statistics below FDR 0.05 for at least one time point and model were plotted for **(A)** cell-type proportions in the spleen, **(B)** cytokines and **(C)** autoantibodies in the plasma. Entities in each plot were sorted using the complete hierarchical method and Euclidean distance. The color scale represents the model statistic limited to an absolute value of 10. The differential entities were classified as early (circles) or late (triangles).

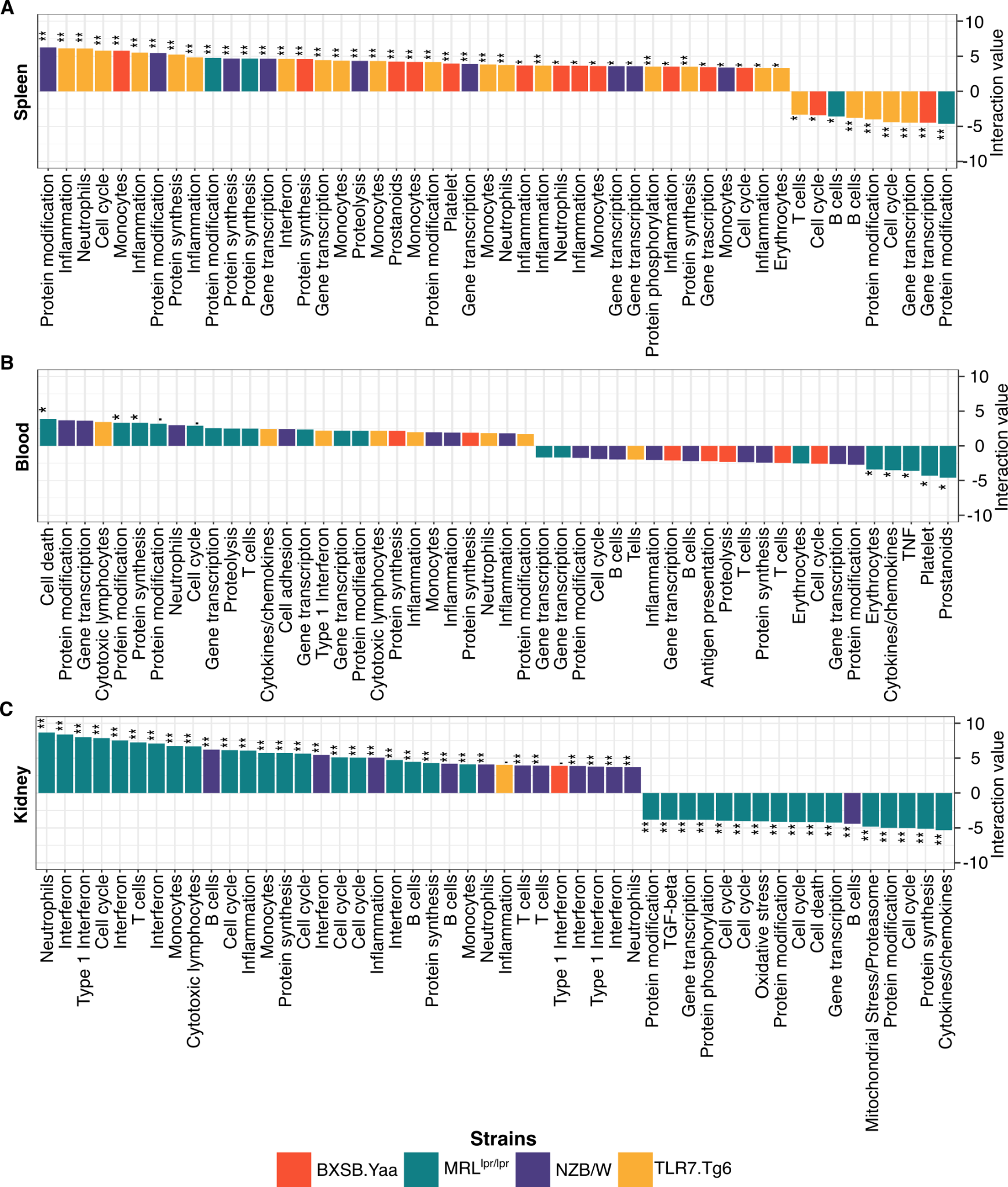
**Fig. S4: Most significant transcriptome module functions per biological sample.**

The figure displays the top 50 highest significant module interaction term values for each tissue: **(A)** spleen, **(B)** blood and **(C)** kidney. The modules were ranked based on the interaction term value and colored according to the mouse model: yellow (Tlr7.Tg6), purple (NZB/W), green (MRL*^lpr/lpr^*), and red (BXSB.*Yaa*). Modules without defined molecular functions were excluded from the plot. Interaction term significant values are indicated as follows: . FDR < 0.1, * FDR < 0.05, ** FDR < 0.01.

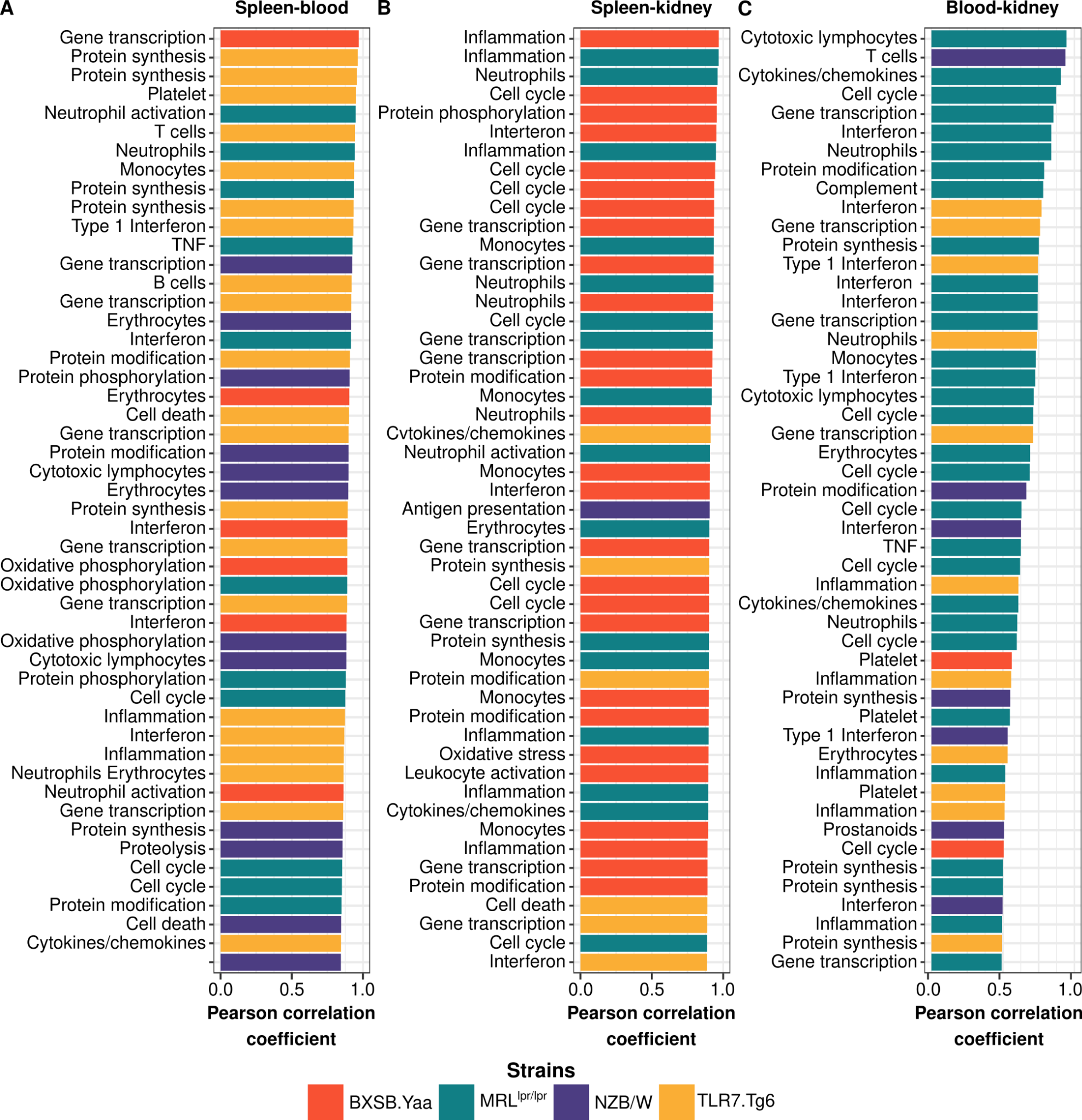
**Fig. S5: Most correlated transcriptome module functions per pair of mouse biological samples.**

The figure displays the top 50 highest correlated modules for each pair of biological samples: **(A)** spleen-blood, **(B)** spleen-kidney and **(C)** blood-kidney. Modules were ranked by Pearson correlation coefficients and colored according to the mouse model: yellow (Tlr7.Tg6), purple (NZB/W), green (MRL*^lpr/lpr^*), and red (BXSB.*Yaa*). Modules without defined molecular functions were excluded from the plot.

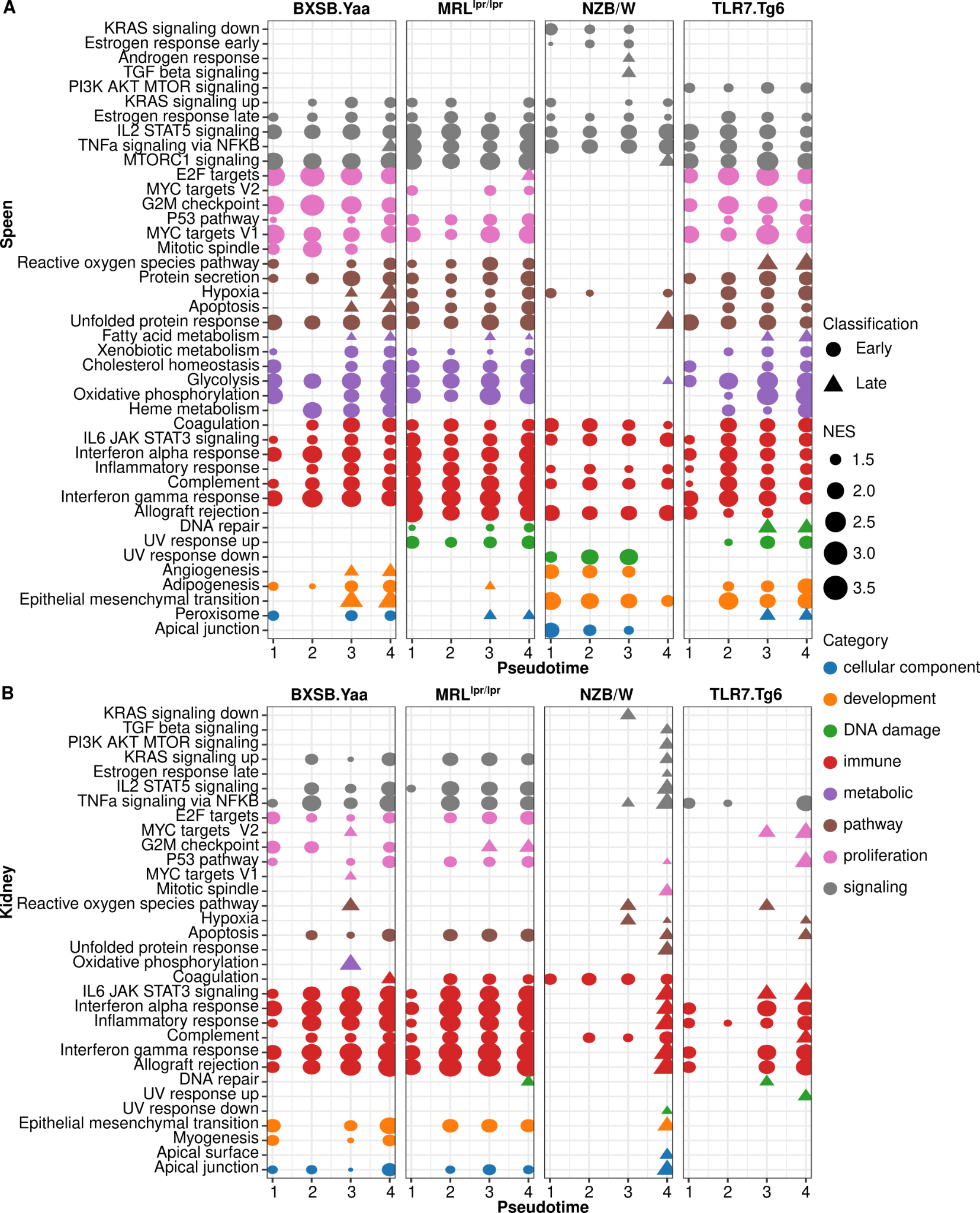

**Fig. S6: Transcriptome GSEA time point analysis by mouse model.**

GSEA was performed on MgSigDB hallmark gene sets, **(A)** spleen and **(B)** kidney normalized positive enrichment scores (NES) below FDR 0.05 for at least one time point and mouse model were plotted, grouped and colored by gene set categories. Hallmark gene sets were classified as early (circles) or late (triangles).

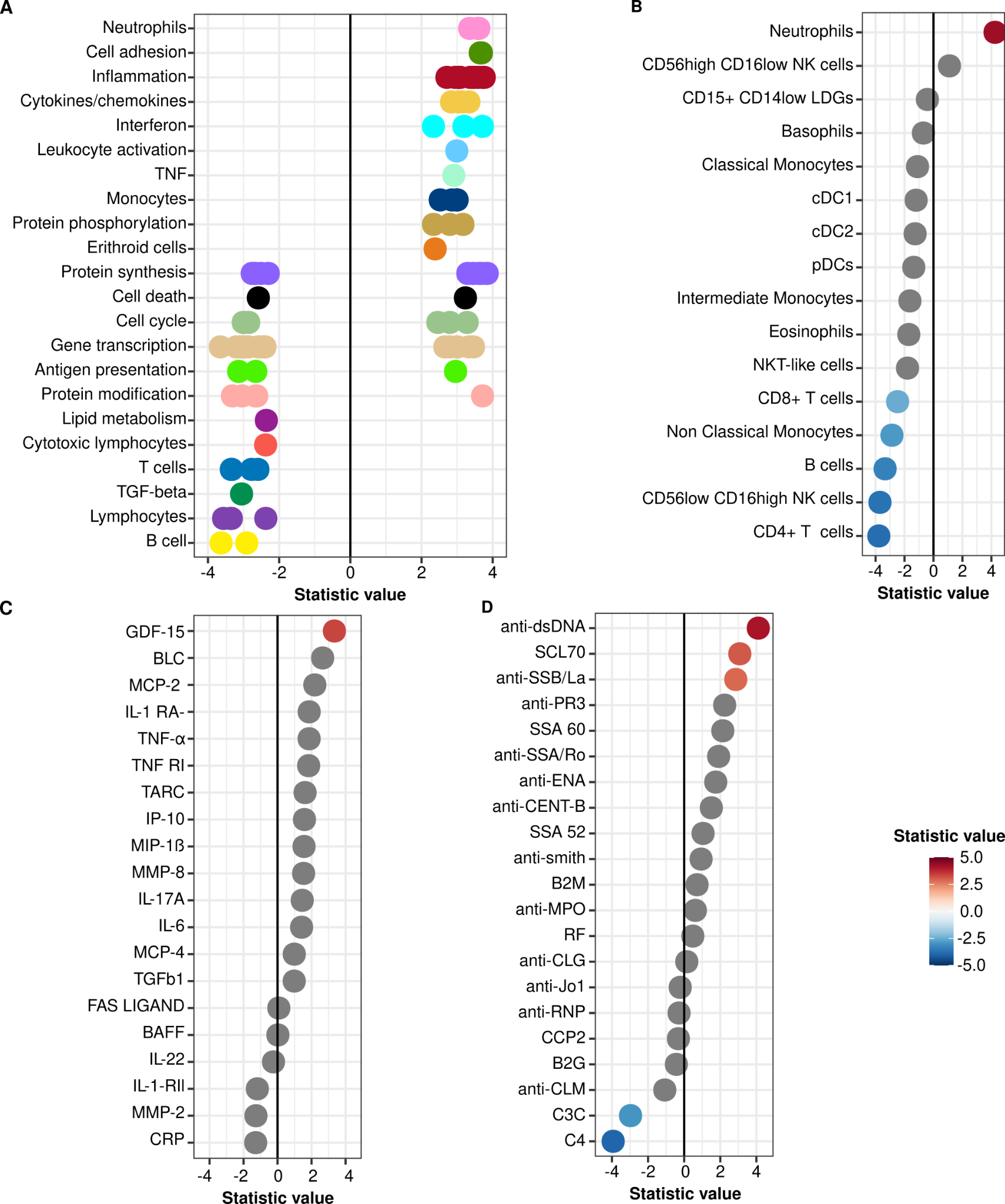

**Fig. S7: Association of molecular features with the human SLEDAI-2K activity index.**

Significant features (FDR < 0.05) are grouped and colored by **(A)** functional module definition for blood transcriptome modules and ranked by regression coefficient average. **(B)** Blood cell populations, **(C)** cytokines and **(D)** autoantibodies are ranked by regression coefficients and colored in red-blue color-scale if significant (FDR < 0.05).

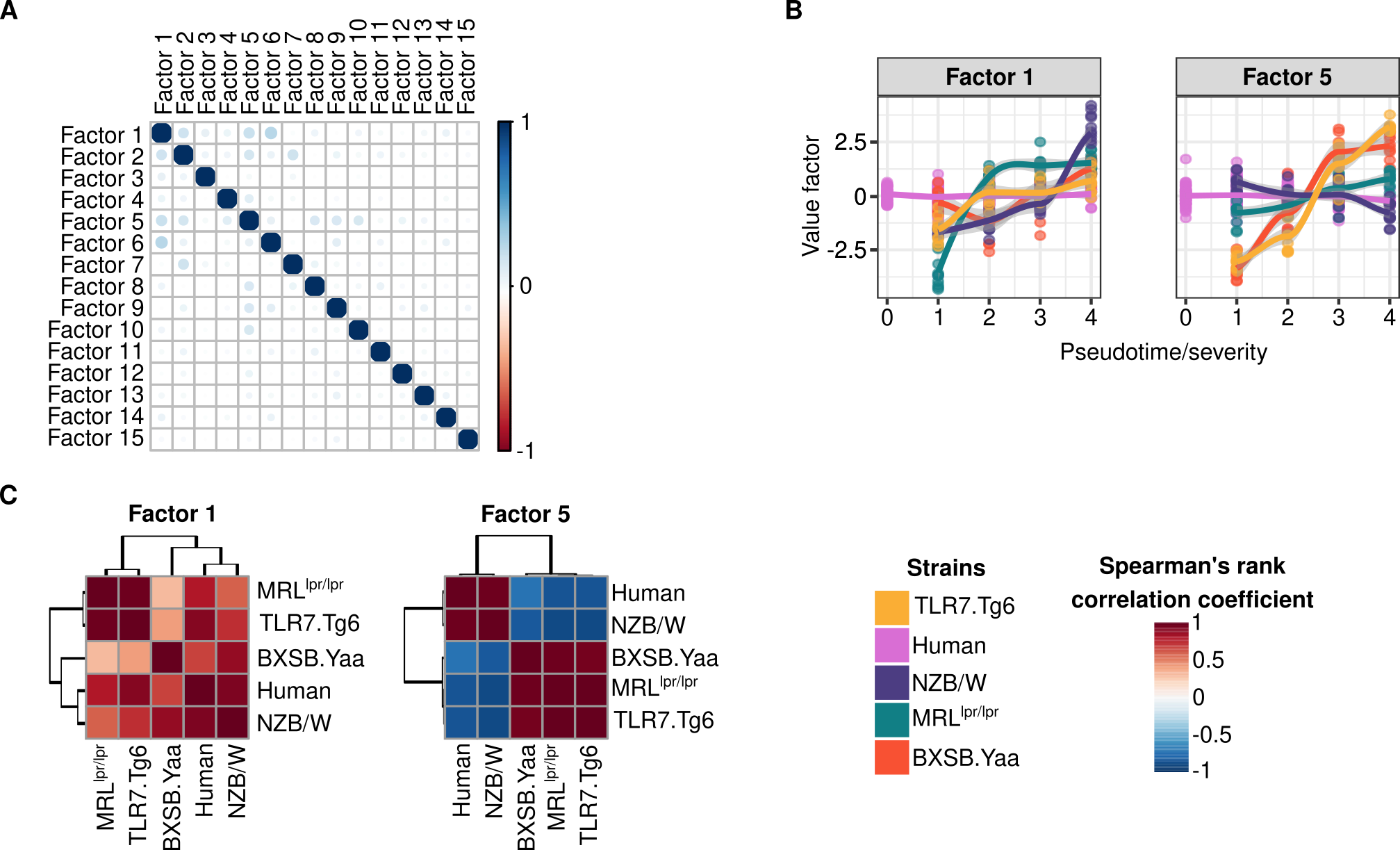

**Fig. S8: Factor correlations and relationships for Factors without significant trends in humans.**

**(A)** Correlations between factor values are shown**. (B)** Trend along severity is shown for each factor per group. Factor values per severity and group are summarized by means of loess regression. **(C)** Factor value correlations between human and mouse models are depicted. Mouse models are color-coded as follows: yellow (Tlr7.Tg6), purple (NZB/W), green (MRL*^lpr/lpr^*), and red (BXSB.*Yaa*). Human results are colored pink.

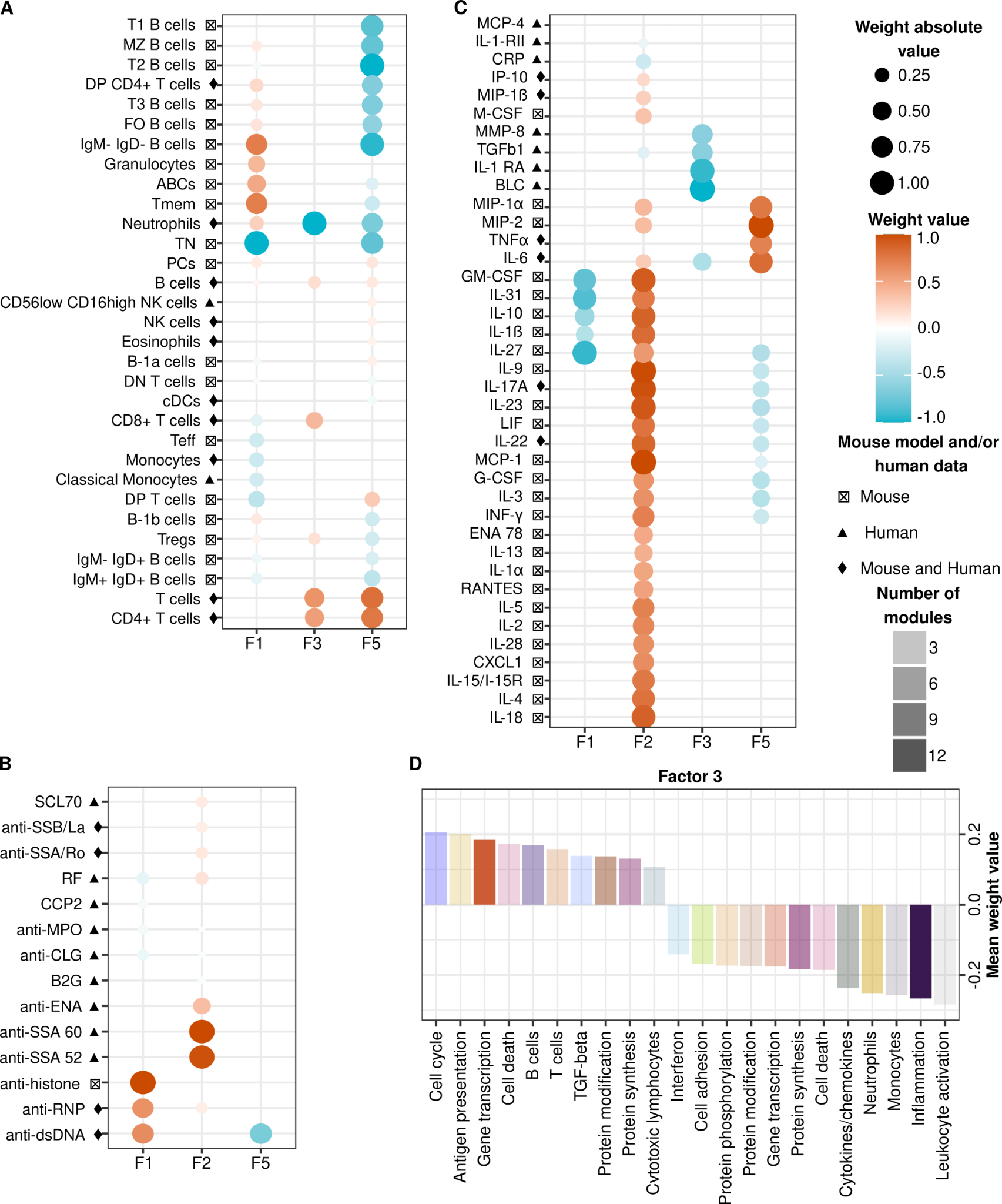

**Fig. S9: Factors are contributed by immune mediated signatures previously defined in SLE pathogenesis.**

Representative positive factor features were selected above 95th percentile and negatives under 5th percentile. Weight absolute values are shown per molecular entity (rows) and Factor (columns) for **(A)** cell populations, **(B)** autoantibodies and **(C)** cytokines. **(D)** Transcriptional modules shared between mouse spleen and human blood contributing to Factor 3 are shown ranked by mean weight value and colored by functionality.

|  | **CTRL** | **SLE** |
| --- | --- | --- |
| **Total (n)** | 564 | 379 |
| **Age (X±SD)** | 47.0 ± 13.1 | 46.1 ± 13.8 |
| **Female (%)** | 79,4 | 93,1 |
| **SLEDAI (X±SD)** | NA | 5.94 ± 5.27 |
| *No activity (n)* | NA | 44 |
| *Mild (n)* | NA | 171 |
| *Moderate (n)* | NA | 100 |
| *High (n)* | NA | 52 |
| *Very high (n)* | NA | 11 |
| **Disease Duration *years* (X±SD)** | NA | 14.6 ± 9.7 |
| **Biopsy proven Nephritis** | 0 | 93 |
| **Medication (%)** | | |
| **Steroids** | NA | 49.6 |
| **Antimalarials** | NA | 70.7 |
| **Immunosuppressants** | NA | 34.8 |
| **Biologics** | NA | 0 |
| **Off-Treatment** | NA | 14.0 |
| **Molecular Information** | | |
| **Transcriptome (n)** | 497 | 342 |
| **Cytokines (n)** | 317 | 262 |
| **Autoantibodies (n)** | 487 | 307 |
| **Flow Cytometry (n)** | 535 | 358 |

**Table S1: Human cohort description.**

|  |  | **BXSB.*Yaa*** | **BSXB** | ***MRL^lpr/lpr^*** | **MRL/J** | **NZB/W** | **NZB** | **Tlr7.Tg6** | **C57/Bl6** |
| --- | --- | --- | --- | --- | --- | --- | --- | --- | --- |
| **RNA-Seq**  **Blood** | *PTime 1* | 8 | 4 | 7 | 2 | 10 | 5 | 6 | 3 |
|  | *PTime 2* | 5 | 2 | 5 | 3 | 10 | 5 | 9 | 5 |
|  | *PTime 3* | 8 | 3 | 6 | 3 | 8 | 5 | 9 | 5 |
|  | *PTime 4* | 8 | 4 | 5 | 3 | 9 | 5 | 8 | 5 |
| **RNA-Seq**  **Kidney** | *PTime 1* | 10 | 5 | 10 | 5 | 9 | 5 | 10 | 4 |
|  | *PTime 2* | 10 | 5 | 9 | 5 | 10 | 5 | 10 | 5 |
|  | *PTime 3* | 10 | 5 | 10 | 5 | 8 | 5 | 10 | 5 |
|  | *PTime 4* | 9 | 5 | 11 | 5 | 12 | 6 | 9 | 7 |
| **RNA-Seq**  **Spleen** | *PTime 1* | 10 | 5 | 10 | 4 | 10 | 5 | 9 | 5 |
|  | *PTime 2* | 10 | 5 | 8 | 5 | 10 | 5 | 10 | 4 |
|  | *PTime 3* | 10 | 5 | 10 | 5 | 9 | 5 | 10 | 5 |
|  | *PTime 4* | 9 | 4 | 11 | 5 | 12 | 6 | 10 | 7 |
| **Cytokines** | *PTime 1* | 8 | 4 | 7 | 2 | 10 | 5 | 6 | 3 |
|  | *PTime 2* | 5 | 2 | 5 | 3 | 10 | 5 | 9 | 5 |
|  | *PTime 3* | 8 | 3 | 6 | 3 | 8 | 5 | 9 | 5 |
|  | *PTime 4* | 8 | 4 | 5 | 3 | 9 | 5 | 8 | 5 |
| **AutoAbs** | *PTime 1* | 8 | 4 | 7 | 2 | 10 | 5 | 6 | 3 |
|  | *PTime 2* | 5 | 1 | 5 | 3 | 10 | 5 | 9 | 5 |
|  | *PTime 3* | 8 | 3 | 6 | 3 | 8 | 5 | 9 | 5 |
|  | *PTime 4* | 8 | 4 | 5 | 3 | 9 | 5 | 8 | 5 |
| **Flow Cytometry Spleen** | *PTime 1* | 10 | 5 | 5 | 4 | 10 | 5 | 9 | 5 |
|  | *PTime 2* | 10 | 5 | 8 | 5 | 10 | 5 | 10 | 4 |
|  | *PTime 3* | 10 | 5 | 10 | 5 | 9 | 5 | 10 | 5 |
|  | *PTime 4* | 9 | 4 | 11 | 5 | 12 | 6 | 10 | 5 |

**Table S2: Mouse models cohort description.** Number of individuals per pseudotime (PTime), molecular information and mouse model are depicted.

| Marker | Fluorochrome | Clone | Isotype | Conc ug/ul | ug for staining | ul for staining | Ref. number | Size | Brand |
| --- | --- | --- | --- | --- | --- | --- | --- | --- | --- |
| CD11c | APC | HL3 | hamster IgG1, l2 | 0.2 | 0.2 | 1 | 561119 | 25ug | BD |
| CD49b | APC | DX5 | rat IgM, k | 0.2 | 0.2 | 1 | 560628 | 50ug | BD |
| CD8a | APC-Cy7 | 53-6.7 | rat IgG2a, k | 0.2 | 0.2 | 1 | 561967 | 25ug | BD |
| Gr-1 | APC-Cy7 | RB6-8C5 | rat IgG2b, k | 0.2 | 0.2 | 1 | 557661 | 100ug | BD |
| CD138 | BV510 | 281-2 | rat IgG2a, k | 0.2 | 0.2 | 1 | 563192 | 50ug | BD |
| Zombie Violet vital dye | eF450 | - | - | - | - | 0.5 | 65-0863-14 | 100 tests | ebioscience |
| IgM | eF660 | II/41 | rat IgG2a, k | 0.2 | 0.25 | 1.25 | 50-5790 | 25ug | ebioscience |
| IgD | eF780 | 11-26 | rat IgG2a, k | 0.2 | 0.125 | 0.63 | 47-5993-80 | 25ug | ebioscience |
| CD5 | FITC | 53-7.3 | rat IgG2a, k | 0.5 | 0.5 | 1 | 553020 | 100ug | BD |
| CD115 | PE | T38-320 | rat IgG1, k | 0.2 | 0.1 | 0.5 | 565249 | 100ug | BD |
| CD25 | PE | PC61 | rat IgG1, l | 0.2 | 0.2 | 1 | 561065 | 25ug | BD |
| CD138 | PE | 281-2 | rat IgG2a, k | 0.2 | 0.1 | 0.5 | 561070 | 25ug | BD |
| CD23 | PE | B3B4 | rat IgG2a, k | 0.2 | 0.125 | 0.63 | 12-0232-81 | 50ug | ebioscience |
| CD44 | PE-Cy7 | IM7 | rat IgG2b, k | 0.2 | 0.25 | 1.25 | 103029 | 25ug | Biolegend |
| CD93 | PE-Cy7 | AA4.1 | rat IgG2b, k | 0.2 | 0.5 | 2.5 | 25-5892-81 | 50ug | ebioscience |
| SiglecH | PE-Cy7 | eBio440c | rat IgG2b, k | 0.2 | 0.25 | 1.25 | 25-0333-80 | 25ug | ebioscience |
| CD21 | PerCP-Cy5.5 | 7e9 | rat IgG2a, k | 0.2 | 0.25 | 1.25 | 123415 | 25ug | Biolegend |
| CD62L | PerCP-Cy5.5 | MEL-14 | rat IgG2a, k | 0.2 | 0.25 | 1.25 | 104431 | 25ug | Biolegend |
| CD11b | PerCP-Cy5.5 | M1/70 | rat IgG2b, k | 0.2 | 0.1 | 0.5 | 561114 | 25ug | BD |
| B220 | V500 | RA3-6B2 | rat IgG2a, k | 0.2 | 0.2 | 1 | 561227 | 25ug | BD |
| CD4 | V500 | RM4-5 | rat IgG2a, k | 0.2 | 0.2 | 1 | 560783 | 25ug | BD |

**Table S3: Spleen flow-cytometry antibody information.** Technical information for the antibody mixes used is shown, including: maker, fluorochrome, clone, isotype, concentration, reference number, size and brand (BD states BD Biosciences).
